# Spaceland: Histology-Guided Reconstruction of High-Resolution Whole-Organ 3D Molecular Atlases from Sparse Spatial Transcriptomics

**DOI:** 10.64898/2026.07.23.739686

**Authors:** Fangfang Xu, Zhenfeng Zhuang, Yun Zhu, Bo Ying, Naiqiao Hou, Weiping Lin, Liansheng Wang, Chaoyong Yang, Jia Song

## Abstract

Reconstructing whole organs in three-dimensional molecular detail is a key step toward building virtual organs for modeling tissue organization, disease progression and drug perturbation responses. However, high-resolution whole-organ spatial transcriptomic profiling remains impractical, forcing a trade-off between reconstruction fidelity and sampling density. Here, we introduce Spaceland, a morphology-guided framework that reconstructs continuous, high-resolution 3D molecular landscapes from sparsely sampled spatial transcriptomic sections and serial H&E histology. Spaceland formulates this task as learning continuous gene-expression fields within a morphology-informed histological space. It constructs a dense 3D morphological scaffold by optical-flow interpolation of foundation model-derived H&E representations and decodes sparse spot-level transcriptomic measurements onto an 8 μm histology-aligned grid. Across mouse olfactory bulb, mouse hemibrain and spatiotemporal planarian regeneration, Spaceland generalized across platforms, tissue scales and biological contexts. In mouse benchmarks, Spaceland bridged 400 μm molecular gaps, resolved sub-spot organization, outperformed ST-based interpolation and 2D H&E-based prediction methods, and remained robust with 160-320 μm H&E intervals. In planarian regeneration, it enabled time-resolved whole-organism analysis from only four Visium sections per stage, revealing dynamic neoblast-neural spatial remodeling. Together, Spaceland shifts 3D molecular atlas construction from exhaustive experimental sampling toward data-driven virtual tissue and organ modeling, providing a scalable route to whole-organ molecular reconstruction.

## Introduction

Resolving the three-dimensional molecular architecture of intact organs is essential for understanding how cellular states, tissue structures and molecular programs are organized across complex biological systems^1–4^. Emerging volumetric profiling technologies can map selected RNA species across intact organs^5,6^ or measure highly multiplexed gene expression within three-dimensional tissue blocks^7,8^, but remain constrained by a trade-off between tissue scale and molecular throughput^9,14^.

Reconstruction from serial two-dimensional spatial transcriptomics (ST) sections offers a scalable alternative, but shifts the bottleneck to the axial sampling density and in-plane molecular resolution required for faithful volumetric inference^1,10–13^. Densely sampled, high-resolution ST sections improve reconstruction fidelity but remain costly and time-consuming^3,4,9^. ST-based imputation and interpolation methods can partially alleviate this burden by inferring missing transcriptomic signals from sparse or low-resolution measurements^1,14–16^. However, because they rely primarily on transcriptomic continuity across aligned sections, these methods become increasingly underconstrained as inter-section spacing increases or in-plane resolution decreases, limiting their ability to recover fine tissue-aligned molecular structures and local expression boundaries^14,17^.

Morphology-assisted reconstruction provides an opportunity to use dense structural information to constrain molecular inference within and between sparsely sampled ST sections. Existing approaches based on three-dimensional structural modalities such as microCT can provide volumetric priors, but are limited to low-resolution transcriptomic volumes and introduce additional imaging burden outside routine ST workflows^17^. By contrast, serial H&E histology is routinely acquired alongside ST, inexpensive to scale and captures high-resolution anatomical continuity across tissue volumes, making it a natural dense morphological prior for volumetric molecular reconstruction. Yet current histology-guided transcriptomic prediction remains largely two-dimensional^18–20^, whereas existing ST-based interpolation methods do not fully exploit dense histological continuity. This leaves no scalable framework that jointly uses sparse molecular measurements and serial histology to infer continuous high-resolution three-dimensional molecular fields.

Here, we present Spaceland, a morphology-guided computational framework for reconstructing continuous, high-resolution three-dimensional molecular landscapes from sparsely sampled ST sections and serial H&E histology. Spaceland learns continuous gene-expression fields within a histology-informed three-dimensional morphological space by first constructing a dense H&E-derived morphological scaffold through foundation model feature extraction and optical flow-based interpolation, and then linking this scaffold to sparse spot-level ST measurements using a tailored multiple-instance learning decoder. This design jointly addresses axial anisotropy from sparse molecular sampling and limited in-plane resolution from spot-based measurements, enabling three-dimensional gene-expression inference on a near-isotropic, histology-aligned 8 μm sub-spot grid.

We validated Spaceland across spatial transcriptomic platforms, tissue scales and biological contexts. In a mouse olfactory bulb benchmark, Spaceland reconstructed high-resolution molecular landscapes from only eight 50 μm-spot ST sections separated by up to 400 μm, using serial H&E histology as a dense morphological scaffold. This enabled Spaceland to mitigate the axial anisotropy from sparse molecular sampling, recover fine-scale spatial organization beyond native spot resolution, outperform representative reconstruction and interpolation approaches, and remain robust when H&E sampling intervals were increased to 160-320 μm. These advantages further extended to coarser-resolution whole-brain reconstruction. Finally, in regenerating planarians, Spaceland enabled volumetric analysis of dynamic molecular organization, revealing temporally coordinated remodeling of stem-cell and neural-associated spatial relationships during whole-organism regeneration. Together, Spaceland provides a scalable framework for building continuous, high-resolution three-dimensional molecular tissue models, offering a step toward virtual organs that capture tissue architecture, molecular organization and dynamic biological processes.

## Results

### Overview of Spaceland

Spaceland is a histology-guided framework for reconstructing continuous three-dimensional molecular expression fields from sparsely sampled spot-level ST measurements and serial H&E histology (**Fig. 1a**). The central idea of Spaceland is to treat serial histology as a dense, queryable morphological coordinate system and sparse ST section as molecular constraints within this space. Spaceland achieves this by formulating scalable volumetric reconstruction as two coupled inference problems: first, constructing a continuous histological manifold from incomplete serial H&E sections; and second, learning a molecular field over this manifold from sparse spot-level transcriptomic constraints. This design enables molecular reconstruction across both profiled and unprofiled tissue regions, extending beyond the native spot resolution and axial sampling density of ST data.

**Fig. 1.**
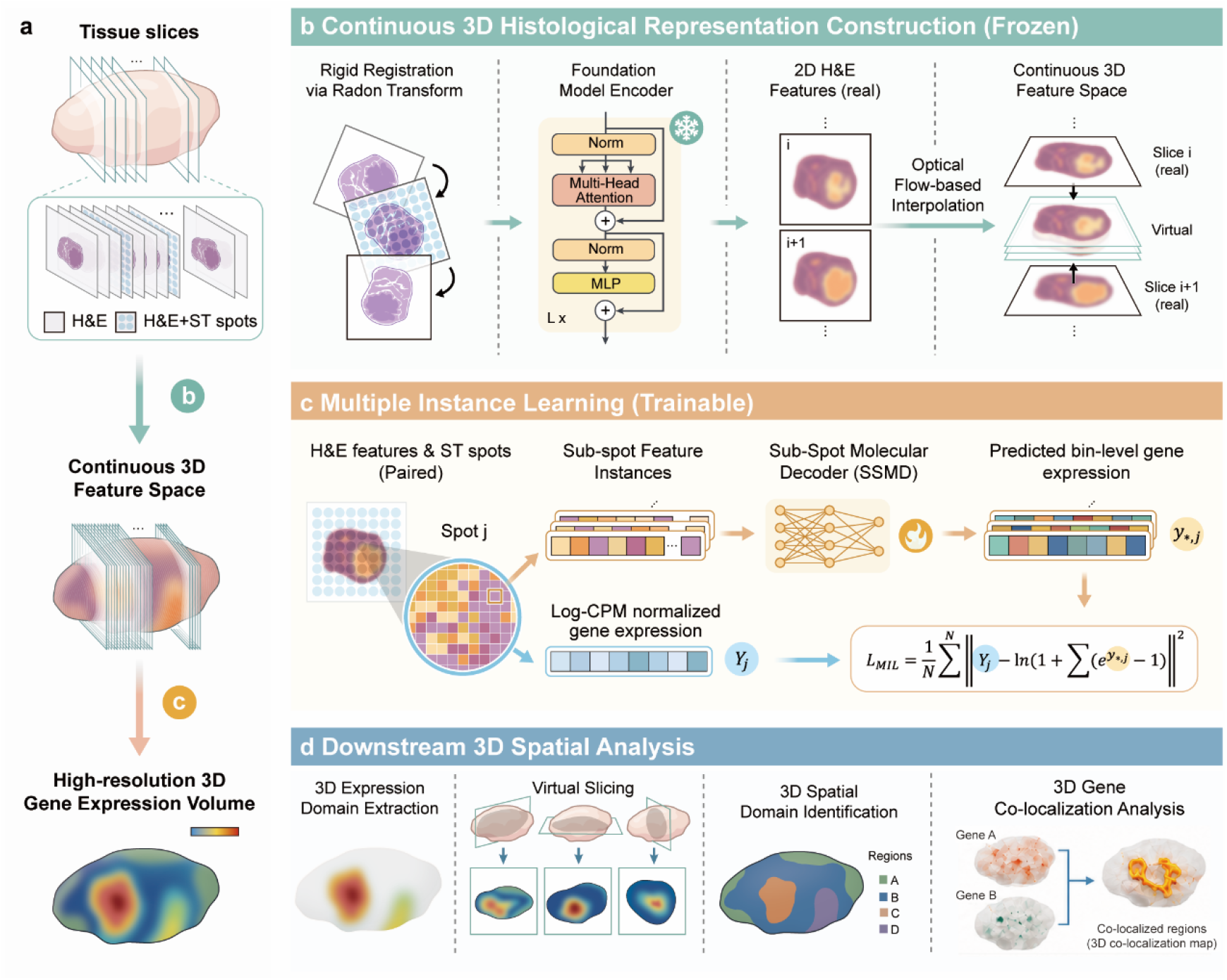
Overview of Spaceland workflow. **a,** Schematic overview of the Spaceland framework. Integrating sparse spatial transcriptomics (ST) data with serial H&E images, Spaceland constructs a near-isotropic 3D morphological representation to generate a high-resolution, dense 3D gene expression volume via multiple instance learning (MIL). **b,** Continuous 3D histological representation construction. Serial H&E images are aligned via Radon transform-based rigid registration. A frozen foundation model extracts 2D histological features, followed by optical flow-based interpolation to infer virtual slices within unmeasured Z-axis gaps, yielding a continuous 3D feature space. **c,** MIL architecture. High-resolution H&E features within an ST spot serve as “instances”, paired with Log-CPM normalized gene expression as “bag labels”. A trainable Sub-Spot Molecular Decoder (SSMD) predicts bin-level (8 μm) expression, optimized end-to-end via a customized loss function that aggregates sub-spot predictions against observed spot-level expression.**d,** Downstream 3D spatial analysis. The Spaceland pipeline supports downstream 3D spatial analyses, including 3D expression-domain extraction from the reconstructed high-resolution molecular volume, orthogonal virtual slicing along XY, YZ and XZ planes, semi-supervised 3D spatial-domain identification, and high-resolution 3D gene co-localization analysis for mapping volumetric co-expression regions.

The first inference problem is to construct a continuous 3D histological scaffold in which molecular expression can be decoded (**Fig. 1b**). Serial H&E sections were first assembled into a common histological coordinate system using fast Radon transform-based rigid registration^21^. To enable scalable processing of large section stacks, global rotation and translation parameters were estimated from Gaussian-smoothed, downsampled whole-tissue images and then applied to the corresponding high-resolution sections. This coarse-to-full-resolution strategy avoided computationally intensive full-resolution optimization while preserving the spatial alignment required for downstream 3D reconstruction. Each registered section is then encoded with CONCH^22^, a large-scale pretrained vision-language pathology foundation model, to obtain compact histological feature representations that retain morphology-relevant information while reducing the computational and storage burden of downstream reconstruction. Moreover, serial histology itself is not always complete: cost-saving interval staining, missing sections, section damage and deformation create gaps in the morphological record that conventional section-wise prediction cannot bridge. To move beyond a discrete stack of stained sections toward continuous morphological modeling, Spaceland applies optical flow-based nonlinear interpolation^23,24^ between adjacent H&E sections to infer virtual histological features at unstained intermediate tissue locations. Together, these steps yield an efficient, dense 3D histological scaffold for molecular reconstruction.

The second inference problem is to learn a molecular expression field over this scaffold through a 3D morphology-to-expression decoder trained with sparse spot-level ST measurements (**Fig. 1c**). Because ST provides spot-level measurements whereas reconstruction is performed at sub-spot resolution, each measured spot is represented as a bag of high-resolution H&E-derived instances sampled from the scaffold, with the observed spot-level expression profile supervising the bag. Within this multiple-instance learning formulation, the Sub-Spot Molecular Decoder predicts gene expression at 8 μm bin resolution throughout the 3D histological space. Sub-spot predictions are first aggregated in linear expression space to preserve the additive nature of molecular counts, and the aggregated and observed spot-level profiles are then compared using a log-stabilized multiple-instance learning loss to reduce sensitivity to extreme expression values. Once trained on sparse ST sections, the decoder is deployed across the full H&E-derived feature field to reconstruct continuous high-resolution 3D gene-expression fields beyond both the native spot resolution and the axial sampling density of ST data.

The resulting molecular volumes enable analyses beyond conventional section-based ST profiling **(Fig. 1d)**. By representing gene expression as continuous 3D fields within an anatomically coherent scaffold, Spaceland supports orthogonal-plane virtual sectioning, volumetric delineation of gene-specific expression domains, high-resolution 3D gene co-localization analysis and semisupervised identification of 3D spatial domains. This transformation from sparse molecular measurements to organ-scale 3D molecular representations provides a computational substrate for interrogating molecular heterogeneity, co-expression organization and spatial architecture across complex tissues.

### Dense histology scaffolds bridge sparse molecular sampling for near-isotropic 3D reconstruction

We first established a densely sectioned mouse olfactory bulb (MOB) benchmark comprising 180 consecutive 10 μm-interval H&E-stained coronal sections to test whether serial histology can bridge large gaps between sparsely profiled molecular sections and support high-resolution three-dimensional gene-expression reconstruction (**Fig. 2a**). To emulate realistic whole-organ atlas settings, only eight evenly spaced sections were profiled by Decoder-seq^25^, leaving approximately 400 μm gaps between adjacent molecular measurements. The intervening serial H&E sections provided a dense morphological scaffold for reconstruction, whereas three additional sections profiled by high-resolution in situ sequencing (ISS)^26^ served as independent orthogonal references for validation.

**Fig. 2.**
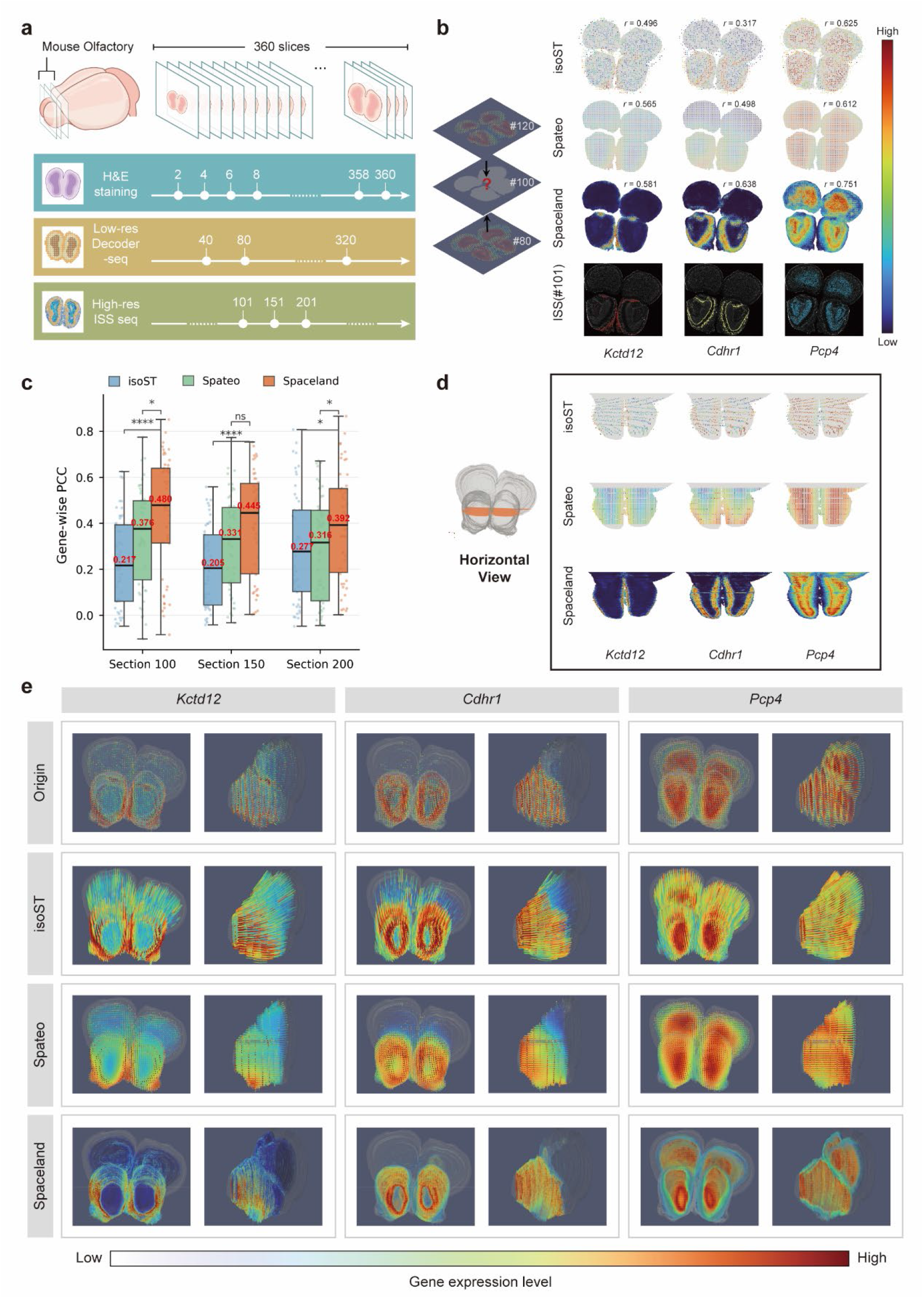
Spaceland reconstructs continuous high-resolution 3D molecular fields from sparsely sampled mouse olfactory bulb sections. **a,** Mouse olfactory bulb benchmark under sparse axial molecular sampling. The dataset comprised 180 H&E-stained coronal sections at 10-μm intervals, eight spot-level Decoder-seq sections separated by approximately 400 μm, and three adjacent high-resolution ISS sections as orthogonal validation references. **b,** Prediction of a molecularly unmeasured section. Expression maps for section 100 predicted by isoST, Spateo and Spaceland are compared with the adjacent ISS section 101 as an orthogonal reference. Spot-level Pearson correlation coefficients are shown above the maps. *Kctd12*, *Cdhr1* and *Pcp4* mark the olfactory nerve, mitral cell and granule cell layers, respectively. Comparisons for sections 150 and 200 are shown in **Extended Data Fig. 1a,b**. **c,** Spot-level reconstruction accuracy on molecularly unmeasured MOB sections. Gene-wise Pearson correlation coefficients (PCCs) were calculated between predictions for sections 100, 150 and 200 and adjacent ISS references from sections 101, 151 and 201, respectively, using the top 50 ISS-derived spatially variable genes ranked by Moran’s I. Each point denotes one gene. Boxes show median and interquartile range; whiskers extend to 1.5× interquartile range. Statistical significance was assessed between methods within each section; ns, not significant; *P < 0.05; ****P < 0.0001. Red numbers indicate median PCC for each method. Comparison with SpatialZ is shown in **Extended Data Fig. 1c**. **d,** Horizontal virtual molecular sections queried from the reconstructed 3D expression fields. Schematics indicate the queried planes. Expression patterns of Kctd12, Cdhr1 and Pcp4 are shown for isoST, Spateo and Spaceland. Sagittal virtual sections are shown in **Extended Data Fig. 1d**. **e,** Three-dimensional renderings of reconstructed gene-expression volumes. Front and side views are shown for the input Decoder-seq sections and the corresponding isoST, Spateo and Spaceland reconstructions.

We next evaluated whether Spaceland could recover molecularly unmeasured sections across these large axial gaps. Using sparse Decoder-seq measurements and the serial H&E scaffold, Spaceland reconstructed gene-expression maps for held-out sections and was benchmarked against representative ST-based reconstruction methods (**Supplementary Table S1**), with adjacent ISS measurements serving as orthogonal references (**Fig. 2b** and **Extended Data Fig. 1a,b**). For marker genes defining major MOB layers^27^, including *Kctd12, Cdhr1* and *Pcp4*, isoST^14^ produced fragmented and spatially discontinuous patterns, whereas Spateo^1^ generated smoother maps with attenuated layer-specific boundaries. In contrast, Spaceland recovered high-resolution expression patterns that closely matched ISS measurements and preserved the laminar organization of MOB marker-gene expression.

This qualitative improvement was supported by quantitative evaluation at matched spot-level resolution. Because isoST and Spateo do not perform sub-spot molecular reconstruction, all methods were compared at the common spot-level scale using gene-wise Pearson correlations for the top 50 spatially variable genes identified from the ISS reference sections. Across sections 100, 150 and 200, Spaceland consistently achieved the highest median PCCs, ranging from 0.392 to 0.480, compared with 0.205–0.277 for isoST and 0.316–0.376 for Spateo (**Fig. 2c**). Although SpatialZ^15^ was originally designed for single-cell-resolution spatial reconstruction and is not directly optimized for spot-level data, an adapted benchmark showed that Spaceland consistently outperformed SpatialZ across all three unmeasured sections, with median PCCs of 0.392–0.480 versus 0.152–0.242 (**Extended Data Fig. 1c**). This common-resolution evaluation provides a fair but conservative assessment of Spaceland, as its native sub-spot predictions were downsampled to the resolution shared by all methods. Beyond these spot-level metrics, Spaceland’s native sub-spot reconstruction further recovered fine laminar structures and sharp molecular boundaries beyond what can be quantified by spot-level correlations alone.

We then evaluated whether the gains observed on held-out sections translated into coherent three-dimensional molecular fields. Reconstructed expression volumes were queried along sagittal and horizontal planes to assess molecular continuity beyond the originally sampled coronal sections (**Extended Data Fig. 1d** and **Fig. 2d**). In these orthogonal views, isoST and Spateo showed pronounced axial artifacts, including fragmented stripe-like signals and over-smoothed band-like patterns with poorly resolved anatomical boundaries. By contrast, Spaceland bridged the 400-µm molecular sampling gaps to produce smooth, anatomically aligned expression gradients across unseen planes while preserving the laminar organization of *Kctd12, Cdhr1* and *Pcp4*. Consistent with the virtual-section analysis, three-dimensional renderings further showed that Spaceland reduced the volumetric fragmentation and banding artifacts apparent in ST-based reconstructions under sparse axial sampling (**Fig. 2e**). Whereas isoST and Spateo remained constrained by the sparse geometry of the profiled molecular sections, Spaceland generated spatially continuous molecular domains that followed the MOB morphology across both front and side views. Together, these results show that Spaceland can use dense serial histology as a morphological scaffold to transform sparse molecular measurements into near-isotropic, high-resolution three-dimensional gene-expression fields.

### Spaceland enables sub-spot molecular decoding with reduced H&E sampling

Having shown that Spaceland can use dense histology to bridge large gaps between sparsely sampled molecular sections, we next asked whether its histology-guided representation could further support high-resolution molecular decoding. Specifically, we investigated whether Spaceland could recover sub-spot gene-expression patterns from H&E morphology and maintain molecular prediction when histological features were interpolated from increasingly sparse H&E section sampling (**Extended Data Fig. 2**).

To test this, we designed an orthogonal validation experiment in which molecularly unmeasured H&E sections were predicted and evaluated against adjacent high-resolution ISS references. Spaceland was trained on eight sparse Decoder-seq sections distributed along the section axis and used to generate 8 μm-resolution predictions for target sections 100, 150 and 200, which were compared with adjacent ISS sections 101, 151 and 201, respectively (detailed training-section indices are provided in **Methods**). This design enabled independent molecular validation of sub-spot predictions while also testing generalization across target sections at different distances from the training data. Because the ISS panel measured a limited set of genes, quantitative analyses focused on the top 50 spatially variable ISS genes selected by Moran’s I.

We first evaluated performance on section 100, the target section farthest from the training data, located approximately 200 μm from the nearest profiled Decoder-seq sections and therefore providing the most stringent test of cross-section generalization. Spaceland outperformed representative H&E-based gene-expression prediction methods, including iStar^28^, scstGCN^29^, iSCALE^30^ and MISO^31^. After aggregation to 8, 16, 32 and 48 μm resolutions, Spaceland achieved the highest gene-wise PCCs across all resolutions, with median PCCs of 0.110, 0.229, 0.402 and 0.486, respectively, compared with 0.100, 0.189, 0.319 and 0.379 for the strongest baseline at each resolution (**Fig. 3a, left**). At 8 μm resolution, Spaceland also achieved high reference-active structural similarity, with a median SSIM of 0.927; although one baseline reached a comparable SSIM, it showed substantially weaker PCC, indicating that Spaceland more consistently balanced expression accuracy and spatial-structure preservation (**Fig. 3a, right)**. Similar advantages were observed on sections 150 and 200 (**Extended Data Fig. 3**). Notably, baseline performance improved on target sections closer to the training data but was substantially weaker on section 100, indicating sensitivity to cross-section variation and training-section proximity. In contrast, Spaceland showed stronger generalization across molecularly unmeasured sections.

**Fig. 3.**
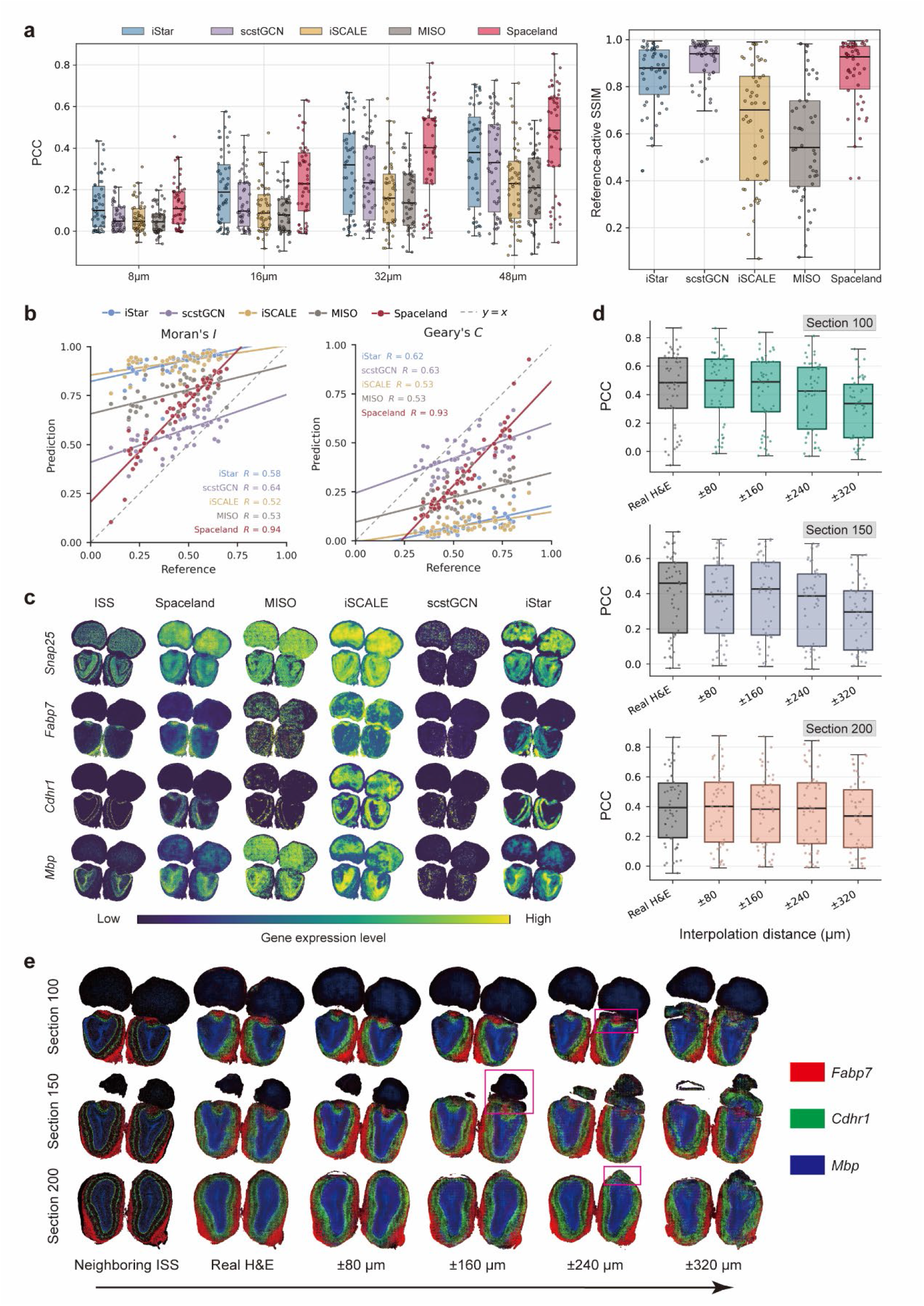
Spaceland predicts high-resolution gene-expression patterns from H&E and preserves performance with interpolated histological features. **a,** Quantitative comparison of Spaceland with representative H&E-based gene-expression prediction baselines on mouse olfactory bulb section 100, evaluated against the adjacent ISS reference section 101. Left, Pearson correlation coefficients (PCCs) for the top 50 spatially variable genes, identified from the ISS reference using Moran’s I, after aggregation to 8, 16, 32 and 48 μm resolutions. Right, 8 μm SSIM computed within reference-active tissue regions for each gene. Each point represents one gene at the indicated resolution. Box plots show the median, interquartile range and 1.5× interquartile-range whiskers. Results for sections 150 and 200 are shown in **Extended Data Fig. 3**. **b,** Concordance of spatial autocorrelation metrics between predicted 8 μm expression maps and the adjacent ISS reference for section 100. Moran’s I and Geary’s C were computed for the same 50 spatially variable genes as in **a**. Each point represents one gene. Solid lines denote method-specific linear regression fits, and dashed grey lines denote the identity line. Pearson correlation coefficients are shown in each panel; corresponding two-sided test P values are provided in **Supplementary Data 1**. Results for sections 150 and 200 are shown in **Extended Data Fig. 4**. **c,** Representative 8 μm expression maps for selected genes with distinct spatial patterns. Predictions for H&E section 100 are shown alongside the adjacent ISS reference section 101. Rows show genes and columns show the ISS reference and predictions from Spaceland and baseline methods. Color scales indicate gene-wise relative expression. Results for sections 150 and 200 are shown in **Supplementary** Fig. 1. **d,** Ablation analysis of optical-flow-interpolated histological features. For target sections 100, 150 and 200, Spaceland was run using either real H&E features from the target section or interpolated H&E features synthesized from flanking sections at increasing axial distances. Predicted expression maps were aggregated to 48 μm and evaluated against the adjacent ISS reference by gene-level PCC. The ±80, ±160, ±240 and ±320 μm labels indicate the axial distances of the flanking H&E sections used for feature interpolation. Each point represents one gene. Box plots show the median, interquartile range and whiskers extending to 1.5× the interquartile range. Results at 8, 16 and 32 μm resolutions are shown in **Supplementary** Fig. 2. **e,** Multi-channel visualization of expression patterns predicted from real and interpolated histological features. Composite maps show *Fabp7* in red, *Cdhr1* in green and *Mbp* in blue for sections 100, 150 and 200. Predictions generated from real H&E features and interpolated features at increasing axial distances are shown alongside adjacent ISS references.

We next asked whether these improvements reflected genuine recovery of spatial organization rather than higher agreement in gene-expression magnitude alone. On section 100, Moran’s I and Geary’s C computed from the 8 μm predictions for this gene set showed strong concordance with the adjacent ISS reference, with Pearson correlation coefficients of 0.94 and 0.93, respectively, substantially exceeding the best-performing baselines (0.64 and 0.63; **Fig. 3b**). The same trend was observed for sections 150 and 200 (**Extended Data Fig. 4**). Representative top spatially variable genes with distinct spatial patterns, including *Snap25, Fabp7, Cdhr1* and *Mbp*, further illustrated these differences: Spaceland recovered laminar and region-specific expression domains that closely matched the ISS reference, whereas baseline methods produced blurred anatomical boundaries, discontinuous local signals or gene-dependent artifacts **(Fig. 3c, Supplementary Fig. 1**).

Finally, we showed that although Spaceland uses H&E histology as a morphological scaffold, it does not require exhaustively dense histological sampling. For sections 100, 150 and 200, Spaceland was run using either real H&E-derived features from the target section or optical-flow-interpolated features synthesized from flanking H&E sections at increasing axial distances. The tolerable interpolation distance depended on local anatomical variability: predictions remained reliable up to ±80 μm in section 150, where morphology changed more rapidly, and up to approximately ±160 μm in sections 100 and 200, where local morphology was more stable (**Fig. 3d**). Multi-channel visualization of *Fabp7, Cdhr1* and *Mbp* confirmed that major laminar expression domains were preserved under these interpolation settings (**Fig. 3e**). These results indicate that Spaceland can use interpolated histological features over 160-320 μm ranges, supporting adaptive H&E sampling strategies that reduce the experimental burden of three-dimensional molecular reconstruction.

Together, these results demonstrate that the histology-guided representation learned by Spaceland supports both sub-spot molecular decoding and histological feature interpolation, enabling high-resolution three-dimensional reconstruction without requiring exhaustive molecular or histological sampling.

### Scalable reconstruction of continuous organ-scale molecular landscapes

To evaluate the scalability of Spaceland to organ-scale tissues with more heterogeneous anatomy and coarser molecular measurements, we applied it to a public adult mouse hemibrain spatial transcriptomic dataset^32^. From the original dataset, we retained 62 coronal sections with paired H&E images and approximately 100 μm-resolution spot-level transcriptomic measurements for analysis (**Fig. 4a** and **Methods**). Because many anterior-posterior (AP) positions contained paired neighboring A/B sections, we trained Spaceland on the smaller B-section subset comprising 29 sections and reserved the A-section subset comprising 33 sections for held-out evaluation across the AP axis (detailed in **Extended Data Fig. 5** and **Supplementary Table S2**).

**Fig. 4.**
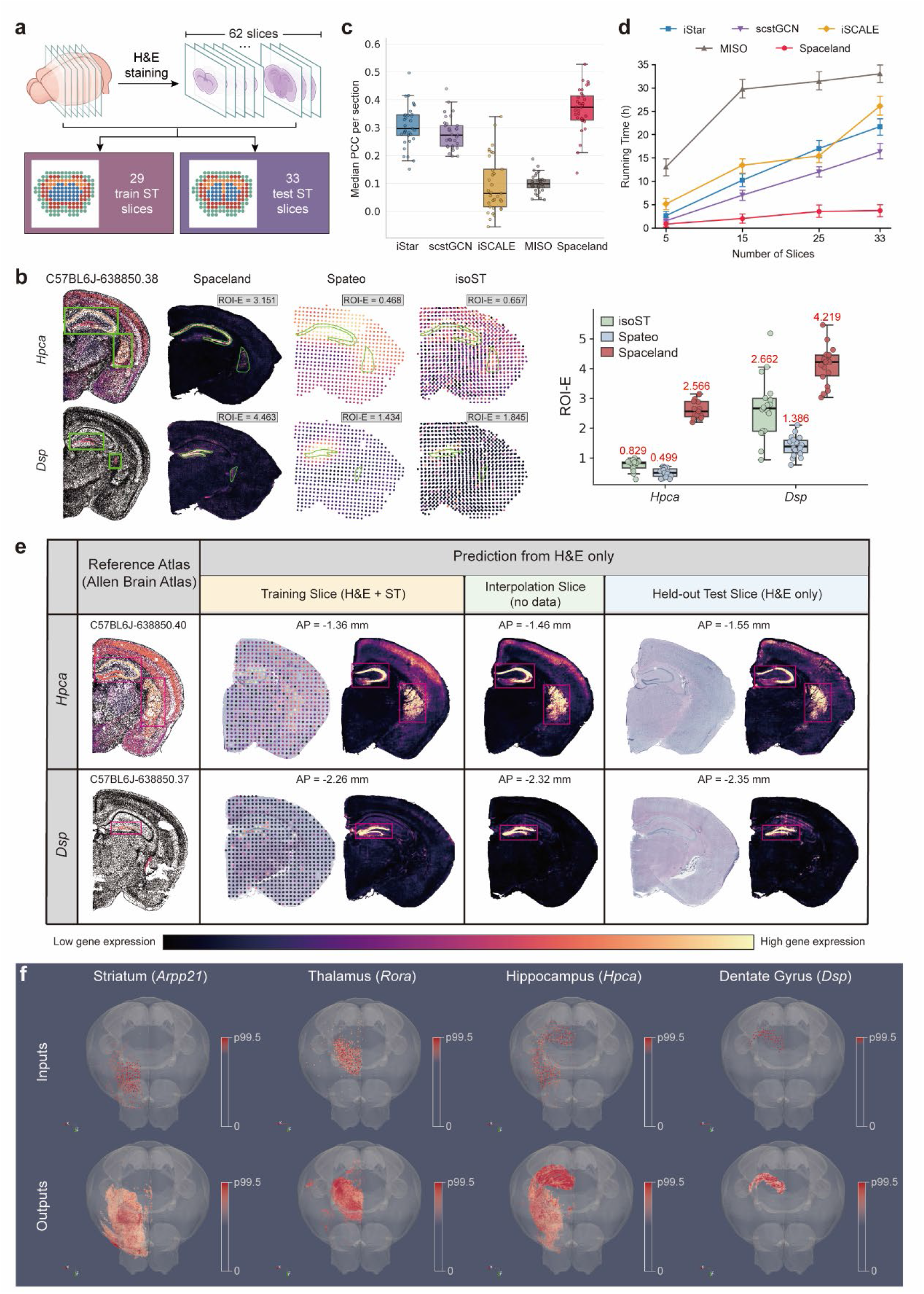
Spaceland reconstructs a scalable high-resolution three-dimensional molecular atlas of the adult mouse hemibrain. **a,** Schematic of the adult mouse hemibrain benchmark. Sixty-two coronal sections were selected from a published adult mouse hemibrain ST dataset along the anterior–posterior (AP) axis, each paired with an H&E image and spot-level transcriptomic measurements with an approximately 100-μm spot diameter. Spaceland was trained on 29 ST sections and evaluated on 33 held-out sections. The spatial distribution of training and held-out sections is shown in **Extended Data Fig. 5**. **b,** ROI-based evaluation of anatomically localized marker-gene enrichment. **Left,** Allen Brain Atlas^33^ -guided ROIs for *Hpca* and *Dsp* were annotated on the H&E image of section 23A (AP = −2.25 mm) and overlaid on predicted expression maps from Spaceland, Spateo and isoST. ROI-E values above each map indicate log2 fold enrichment inside versus outside the annotated ROI within tissue. **Right,** ROI-E quantification across serial coronal sections for *Hpca* (sections 14A–28A; n = 15) and *Dsp* (sections 14A–31A; n = 18). Each point represents one section. Boxes show the median and interquartile range; whiskers extend to 1.5× the interquartile range. Red labels indicate median ROI-E values. **c,** Quantitative evaluation on 33 held-out ST sections. Box plots compare Spaceland with representative H&E-based gene-expression prediction baselines, including iStar, scstGCN, iSCALE and MISO, across the top 300 spatially variable genes. Spaceland sub-spot predictions were aggregated to the spot level before computing gene-wise Pearson correlation coefficients (PCCs). Each point denotes the median PCC across genes for each held-out section. Full per-section PCC distributions and Spearman’s rank correlations are shown in **Extended Data Fig. 6a,b**. **d,** Runtime comparison. End-to-end running time was measured for the complete pipeline required by each method. All algorithms were run on the same hardware, with hardware configuration provided in Methods. Each method was run independently three times for each setting; points and error bars indicate mean ± s.d. **e,** Anatomically resolved expression maps across training, virtual and held-out sections. Columns show approximately matched Allen Brain Atlas MERFISH references^33^ and Spaceland predictions on ST-supervised training sections, virtual interpolated sections without directly acquired H&E or ST measurements, and held-out ST sections. Representative training–held-out pairs are 17B/18A and 23B/24A, with 17B and 23B used for ST supervision and 18A and 24A reserved for evaluation. *Hpca* and *Dsp* show anatomically localized expression patterns across the three section types. **f,** High-resolution three-dimensional reconstruction of brain-region-specific expression domains. Marker genes are shown for the striatum (*Arpp21*), thalamus (*Rora*), hippocampus (*Hpca*) and dentate gyrus (*Dsp*). Sparse input ST measurements with discontinuous axial sampling are shown above Spaceland-reconstructed volumetric expression domains. Color bars indicate normalized expression intensity, scaled from 0 to the 99.5th percentile.

We first compared Spaceland with ST-based 3D reconstruction methods, including Spateo and isoST, in recovering anatomically localized expression structures. Using Allen Brain Atlas-guided gene-positive regions of interest (ROIs) as external anatomical priors, we quantified ROI enrichment (ROI-E, **Supplementary Note 1**). Spaceland generated more spatially confined domains for representative markers including *Hpca* and *Dsp* (**Fig. 4b, left**), with the highest median ROI-E across serial coronal sections: 2.566 for Hpca and 4.219 for Dsp, compared with 0.499-0.829 and 1.386-2.662, respectively, for the ST-based methods (**Fig. 4b, right**). Serial-section visualizations of aggregate predicted expression across the evaluation gene panel further illustrated the denser tissue-wide anatomical detail recovered by Spaceland compared with spot-scale ST-based methods (**Extended Data Fig. 6**). These results indicate that Spaceland better resolves fine-scale, anatomically localized molecular enrichment patterns than spot-scale ST interpolation methods.

Having established this structural advantage over ST-based interpolation methods, we next asked whether Spaceland’s morphology-guided reconstruction strategy also improves molecular super-resolution over H&E-based two-dimensional prediction baselines. For quantitative comparison with measured ST profiles, Spaceland predictions were aggregated to the original ST spot level and evaluated across the top 300 spatially variable genes (**Supplementary Note 1**). Spaceland outperformed these H&E-based baselines, achieving a section-wise median gene-level PCC of 0.373 compared with 0.089–0.297 for the baselines (**Fig. 4c**). This advantage was consistent across most AP positions for both PCC and SCC (**Extended Data Fig. 7a,b**), indicating robust morphology-guided reconstruction performance in a larger, lower-resolution and anatomically heterogeneous brain dataset.

Spaceland also showed favorable computational scalability. End-to-end running time, including H&E feature extraction, model training and inference, reached 3.70 ± 1.30 h for 33 sections, compared with 16.30-33.00 h for H&E-based baselines (**Fig. 4d**). Peak GPU memory and system RAM remained moderate at this scale, reaching 1.82 GB and 13.1 GB, respectively (**Extended Data Fig. 7c,d**). These results show that Spaceland extends to brain-scale reconstruction without prohibitive computational overhead.

We further applied Spaceland for brain-scale virtual molecular sectioning across measured and unmeasured AP positions. By decoding H&E-derived or optical-flow-interpolated histology features, Spaceland generated molecular predictions for training sections, held-out test sections and intermediate virtual sections lacking both ST measurements and directly acquired H&E images (**Fig. 4e**). For representative markers including *Hpca* and *Dsp*, predicted expression patterns were anatomically localized across these section types, with *Hpca* enriched in hippocampal regions and *Dsp* enriched in dentate gyrus. These spatial patterns were concordant with MERFISH-based Allen Brain Atlas reference maps^33^ at matched AP positions, supporting the ability of Spaceland to infer anatomically coherent molecular sections beyond the measured ST slices.

Finally, Spaceland reconstructed high-resolution three-dimensional expression volumes across the mouse hemibrain (**Fig. 4f**). Compared with sparse input ST measurements, the reconstructed volumes formed tissue-aligned molecular domains corresponding to major anatomical structures, including *Arpp21* in the striatum, *Rora* in the thalamus, *Hpca* in the hippocampus and *Dsp* in the dentate gyrus. By converting fragmented sparse measurements into continuous three-dimensional molecular representations of major brain structures, Spaceland provides a foundation for organ- scale analysis of tissue organization and anatomical architecture. Together, these results demonstrate that Spaceland enables anatomically coherent and computationally practical organ-scale molecular reconstruction from sparse spatial transcriptomic measurements.

### Whole-organism 3D molecular reconstruction reveals spatiotemporal remodeling during planarian regeneration

To evaluate Spaceland in a dynamic whole-organism setting, we applied it to a time-resolved multimodal dataset of *Schmidtea mediterranea* regeneration. This dataset^34^ captured injury response, neoblast-associated regeneration and coordinated tissue remodeling across six stages: intact, 6 h post-amputation (hpa), 12 hpa, 24 hpa, 3 d post-amputation (dpa) and 7 dpa (**Fig. 5a**). At each stage, serial H&E sections and paired Visium spatial transcriptomic profiles were collected, with only four Visium sections per time point used for training and the remaining Visium sections held out for evaluation (**Extended Data Fig. 8**).

**Fig. 5.**
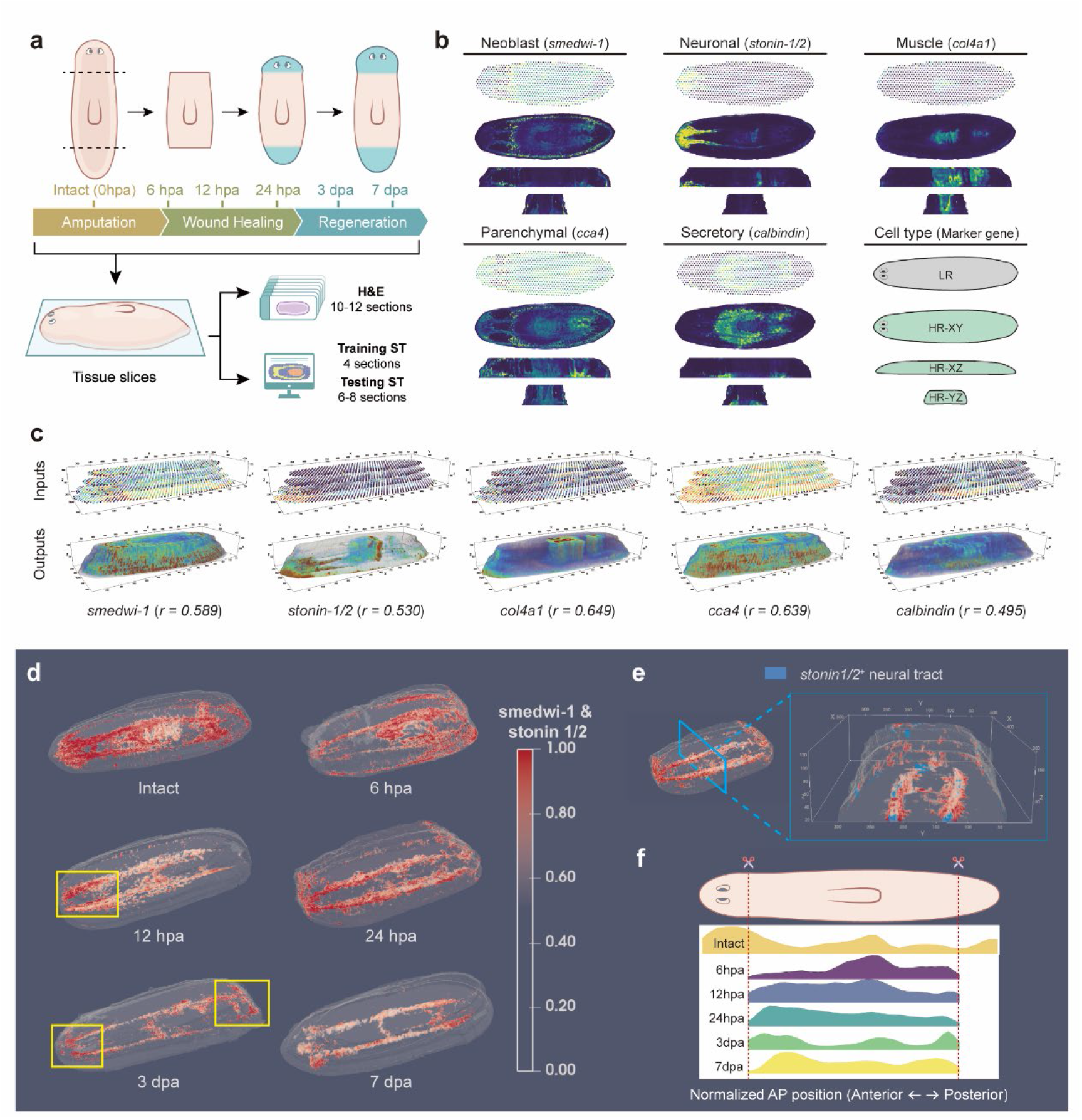
Spaceland reconstructs time-resolved three-dimensional molecular landscapes during planarian regeneration. **a,** Experimental design of the planarian regeneration dataset. Animals were profiled at six stages spanning intact animals, wound healing and regeneration: intact, 6 h post-amputation (hpa), 12 hpa, 24 hpa, 3 d post-amputation (dpa) and 7 dpa. At each stage, serial H&E sections and four paired Visium spatial transcriptomics profiles were collected. Section numbers and train-test splits are provided in **Extended Data Fig. 7**. **b,** Orthogonal virtual molecular sections reconstructed by Spaceland in intact planarians. Low-resolution Visium measurements and Spaceland-reconstructed high-resolution expression maps are shown in the XY, XZ and YZ planes for representative markers of neoblasts (*smedwi-1*), neuronal cells (*stonin-1/2*), muscle (*col4a1*), parenchymal cells (*cca4*) and secretory cells (*calbindin*). **c,** Three-dimensional marker-gene expression volumes. Input Visium measurements are shown in the top row, and Spaceland-reconstructed high-resolution expression volumes are shown in the bottom row. Spot-level Pearson correlation coefficients (*r*) were calculated across all profiled Visium sections after aggregating Spaceland predictions to the Visium spot scale and comparing them with measured Visium expression. **d,** Temporal dynamics of *smedwi-1*–*stonin-1/2* colocalization during regeneration. Three-dimensional maps show normalized local colocalization scores in intact animals and at 6 hpa, 12 hpa, 24 hpa, 3 dpa and 7 dpa. For each gene, expression values were percentile-clipped and Min–Max normalized, and the colocalization score at each three-dimensional spatial point was defined as the pointwise product of normalized *smedwi-1* and *stonin-1/2* expression. Yellow boxes indicate blastema-associated regions with elevated colocalization. Three-dimensional reconstructions of the individual genes are shown in **Extended Data Fig. 10**. **e,** Local three-dimensional view of *smedwi-1*–*stonin-1/2* colocalization at 24 hpa. The colocalization signal is shown together with high *stonin-1/2* expression regions marking the neural tract, highlighting local spatial coupling between neoblast-associated and neuronal programs. **f,** Quantitative profiling of *smedwi-1*–*stonin-1/2* colocalization along the AP axis during regeneration. For each time point, the reconstructed three-dimensional volume was divided into 40 bins along the AP axis. The mean colocalization score within each bin was calculated and Gaussian-smoothed to reduce local noise and generate the ridgeline profiles.

Spaceland reconstructed continuous whole-organism molecular fields from sparse, low-resolution Visium measurements. In intact animals, Spaceland recovered high-resolution expression patterns for representative lineage and tissue markers, including the neoblast marker *smedwi-1*, neuronal marker *stonin-1/2*, muscle marker *col4a1*, parenchymal marker *cca4* and secretory marker *calbindin* **(Fig. 5b)**. Orthogonal virtual sections in the XZ and YZ planes showed continuous spatial expression patterns across the body axis, indicating that the reconstructed molecular fields extended beyond the originally measured XY sections. Three-dimensional renderings further showed that Spaceland transformed sparse Visium measurements into continuous volumetric expression domains while reducing the discrete axial sampling artifacts observed in direct stacking of measured sections **(Fig. 5c)**. After aggregation to the Visium spot scale, reconstructed expression remained consistent with measured Visium profiles across both training and held-out sections, with mean spot-level Pearson correlations of 0.589, 0.530, 0.649, 0.639 and 0.495 for *smedwi-1*, *stonin-1/2*, *col4a1*, *cca4* and *calbindin*, respectively. The reconstructed scaffold also supported volumetric visualization of molecular domains defined from measured Visium sections, yielding spatially coherent annotations of major planarian structures (**Extended Data Fig. 9**).

The time-resolved reconstructions allowed us to examine spatial coupling between neoblast-associated and neural-associated signals during regeneration. Spaceland reconstructed distinct three-dimensional expression patterns for the neoblast marker *smedwi-1* and the neuronal marker *stonin-1/2* across intact and regenerating animals, with reconstructed profiles remaining consistent with measured Visium expression at each time point **(Extended Data Fig. 10)**. We therefore quantified the three-dimensional *smedwi-1*-*stonin-1/2* colocalization to capture the spatial relationship between neoblast-associated activity and neuronal structures. At 24 hpa, local three-dimensional inspection showed that high colocalization was enriched along *stonin-1/2+* neural-tract regions near regenerating tissue rather than diffusely distributed across the animal **(Fig. 5e),** supporting the colocalization score as a local measure of neoblast–neuronal spatial coupling.

Extending this analysis across the full regeneration time course revealed stage-dependent remodeling of *smedwi-1*-*stonin-1/2* spatial coupling **(Fig. 5d)**. In intact animals, high-colocalization regions were broadly aligned with the central nervous system. After amputation, colocalization was reorganized across the body axis. At 6 hpa and 12 hpa, elevated colocalization was preferentially observed near the anterior regenerating region and adjacent neural structures, whereas the posterior wound region showed weaker or more localized signal. From 24 hpa to 3 dpa, high-colocalization regions expanded posteriorly and became more pronounced near the posterior regenerating region. By 7 dpa, both anterior and posterior regions showed enriched colocalization, consistent with progressive restoration of neural-associated spatial organization during regeneration^34^. Quantitative profiling along the AP axis confirmed this temporal shift, with ridgeline profiles showing stage-dependent redistribution of mean colocalization scores across 40 normalized AP bins **(Fig. 5f).**

Together, these analyses demonstrate that Spaceland extends 3D molecular reconstruction from static organ-scale atlases to time-resolved whole-organism molecular atlases. With only sparse Visium supervision at each stage, Spaceland recovered continuous high-resolution molecular fields across the planarian body and enabled volumetric analysis of regeneration-associated spatial coupling between neoblast-associated and neural-associated signals. These results establish serial histology-guided reconstruction as a scalable strategy for resolving dynamic molecular and anatomical remodeling across whole organisms.

## Discussion

In summary, we presented Spaceland, a morphology-guided framework for reconstructing high-resolution three-dimensional molecular tissue landscapes from sparse spatial transcriptomic measurements and serial H&E histology. Rather than treating unmeasured tissue space as a gap to be filled by molecular interpolation between profiled sections, Spaceland formulates reconstruction as learning continuous gene-expression fields within a histology-derived three-dimensional morphological space. In this formulation, sparsely sampled ST sections provide molecular constraints, while serial H&E histology supplies the anatomical continuity that defines the reconstruction space. This changes the basis of volumetric molecular reconstruction from molecular sampling density alone to a joint molecular-morphological representation, improving the scalability of three-dimensional molecular atlas construction.

The results presented here suggest that histology can serve not only as a visual companion to spatial transcriptomics, but also as an active structural constraint for volumetric molecular inference. Across the mouse olfactory bulb and adult mouse hemibrain benchmarks, Spaceland showed that serial H&E information can compensate for two major limitations of section-based ST reconstruction: sparse axial molecular sampling and coarse in-plane spot resolution. Unlike molecular-only three-dimensional ST reconstruction methods, which propagate transcriptomic signals between measured sections and therefore remain constrained by the spacing and native resolution of those molecular measurements, Spaceland anchors sparse ST observations within a dense histology-derived anatomical scaffold to guide molecular inference across unmeasured tissue space. This enables recovery of tissue-aligned molecular patterns across large inter-section gaps, including gaps of up to 400 μm, while supporting sub-spot molecular decoding and orthogonal-plane molecular visualization without exhaustive high-resolution ST profiling. The robustness of Spaceland to reduced H&E sampling, with intervals extended to 160-320 μm depending on local tissue complexity, further indicates that molecular and histological acquisition can be balanced according to tissue structure, desired resolution and available experimental resources. Together, these capabilities provide a practical strategy for scaling three-dimensional molecular atlas construction from benchmark-scale datasets to organ-scale tissues and time-resolved whole-organism systems. In our benchmarks, this enabled reconstruction from as few as eight molecular sections in the olfactory bulb and four sections per stage during planarian regeneration.

As with other spatial reconstruction and imputation approaches^35^, Spaceland’s accuracy varies across genes and is influenced by spatial structure, expression level and technical noise. Its advantage is expected to be strongest for spatially organized genes and gene programs that are reflected, directly or indirectly, in tissue morphology, whereas weakly structured or low-signal genes remain difficult to recover reliably. This scope is consistent with many organ-scale spatial biology applications, which often focus on marker genes, spatial domains, molecular gradients and coordinated gene programs rather than uniform recovery of every transcript. Future extensions incorporating gene-gene co-expression, regulatory priors or transcriptomic foundation models may further expand the range of recoverable genes.

Spaceland also provides a natural computational bridge to emerging volumetric histology, three-dimensional pathology and spatial multi-omics technologies^36,37^. Although the present implementation constructs its morphological scaffold from registered serial H&E sections, future volumetric histology inputs^38^ could reduce registration burden and further improve geometric fidelity. More broadly, the morphology-informed three-dimensional coordinate system defined by Spaceland could serve as a common spatial scaffold for integrating transcriptomic maps with spatial proteomic, multiplexed imaging or emerging spatial epigenomic readouts, allowing morphology and molecular states to be analyzed as spatially coupled features within the same organ-scale tissue volume. Such integrated representations could support virtual tissue atlases in which molecular gradients, anatomical domains, co-localized gene programs and cell-state niches are queried across arbitrary planes and compared across developmental, regenerative, disease or perturbation states. By providing an anatomically grounded and experimentally interpretable route for building three-dimensional molecular tissue models, Spaceland offers a practical step toward data-driven virtual tissues and, ultimately, virtual organs.

## Methods

### Overview of the Spaceland computational framework

Spaceland is a computational framework for reconstructing continuous high-resolution three-dimensional gene-expression volumes by integrating serial H&E-stained histological images with sparsely sampled spatial transcriptomic (ST) measurements. The input consists of a consecutive series of histological sections, denoted as ℐ = *I*_1_, *I*_2_, …, *I_N_*, among which only a sparse subset of sections, indexed by ℳ ⊂ 1,2, …, *N*, is profiled by ST to generate corresponding molecular expression matrices ℰ = *E_m_m∈ℳ__*. For each profiled section *m* ∈ ℳ, the H&E image *I_m_* and the expression matrix *E_m_* are experimentally paired through the ST workflow and spatially aligned during preprocessing. The objective of Spaceland is to infer a dense, histology-aligned three-dimensional molecular expression field across both molecularly measured and unmeasured tissue sections. To this end, Spaceland comprises two coupled components: construction of an near-isotropic three-dimensional morphological representation from serial histology, and multiple-instance-learning-based sub-spot molecular decoding from sparse ST supervision.

### Continuous 3D histological representation construction

#### Data preprocessing

To standardize histological inputs across datasets and experimental batches, all H&E-stained images were resampled to a spatial resolution of 0.5 μm per pixel. At this resolution, each 16 × 16-pixel image tile corresponds to an approximately 8 × 8 μm² tissue region, providing a single-cell-scale spatial unit for subsequent feature extraction and molecular decoding. Stain normalization was then performed using the Macenko method^39^ to reduce non-biological variation in staining intensity and color appearance across sections. Briefly, tissue regions were first identified by threshold-based masking to exclude background pixels. Stain vectors and concentration distributions were then estimated from downsampled tissue images, and each section was normalized with respect to a selected reference section. The resulting normalization transform was applied to the corresponding full-resolution H&E image. For ST data, raw count matrices were normalized using a log-counts-per-million transformation to account for differences in sequencing depth across spots. The normalized expression matrices were used as scale-comparable molecular inputs for subsequent spatial modeling.

#### Global rigid registration based on Radon transform

Sectioning, mounting and imaging can introduce in-plane translation and rotation between consecutive tissue sections, resulting in spatial misalignment along the reconstructed three-dimensional volume. To correct these global section-level offsets, Spaceland adapts the Radon-transform-based rigid registration procedure from CODA^21^ for serial H&E alignment, with modifications tailored to sparse ST-guided volumetric reconstruction. In addition to aligning consecutive H&E images, Spaceland propagates the estimated rigid transformations to the corresponding ST spot coordinates, ensuring that molecular measurements remain spatially registered to the histology-derived three-dimensional scaffold.

For each adjacent section pair, the *i*-th section was used as the fixed image *I_F_*, and the (*i* + 1) -th section was used as the moving image *I_M_* . The in-plane rigid transformation was parameterized by a rotation angle *θ* and a translation vector *t* = (*t_x_, t_y_*)^*T*^. For a point *p* in the moving section, the transformed coordinate *p*′ was defined as

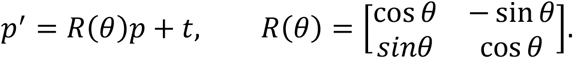

The optimal transformation was estimated by maximizing the normalized cross-correlation between the fixed image and the transformed moving image:

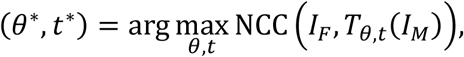

Where *T_θ_*_,*t*_ denotes the rigid transformation applied to the moving image.

Direct joint optimization of rotation and translation on whole-slide images is computationally expensive and can be sensitive to local optima. Following CODA, Spaceland decouples the estimation of rotation and translation using a Radon-transform-based strategy. Before registration, a binary tissue mask was generated for each H&E image to restrict the alignment to tissue-containing regions and reduce the influence of background pixels, coverslip boundaries and scanning artifacts. The Radon transform was then applied to the masked image to project tissue morphology along multiple angles and generate a sinogram. The relative in-plane rotation between adjacent sections was estimated from the angular shift between their Radon-domain representations. After rotation correction, the remaining translational offset was estimated by phase correlation in the spatial domain.

Registration was performed using a coarse-to-fine multiscale strategy. Global alignment was first estimated on downsampled images to obtain a robust initial transformation and was subsequently refined at higher resolutions. For each serial tissue stack, the middle section was selected as the reference section. Adjacent pairwise rigid transformations were estimated along both directions from the middle section and then composed to map all sections into the coordinate frame of this reference. The final transformation for each section was applied consistently to the H&E image and, for molecularly profiled sections, to the corresponding ST spot coordinates. In this way, serial histological images and sparse molecular measurements were mapped into a common coordinate system, providing the aligned multimodal input required for subsequent three-dimensional morphological representation construction and sub-spot molecular decoding.

#### Histological feature extraction

Spaceland uses CONCH^22^, a pathology-specific vision foundation model based on the ViT-B/16 architecture, to extract morphology-aware representations from H&E-stained tissue sections. The CONCH encoder was kept frozen during feature extraction. This design converts each histological section into a dense two-dimensional feature map while avoiding section-specific retraining of the image encoder.

Feature extraction was performed on stain-normalized whole-slide images using a sliding-window strategy. Each image was divided into overlapping windows of 448 × 448 pixels, matching the input size used by CONCH. Adjacent windows were sampled with a stride of 224 pixels, corresponding to 50% overlap in both spatial dimensions. For each window, RGB intensities were normalized using the CLIP mean (0.481,0.458,0.408) and standard deviation (0.269,0.261,0.276) . The ViT-B/16 encoder produced a 28 × 28 grid of patch-level tokens for each window, with one token corresponding to a 16 × 16-pixel image patch and each token represented by a 768-dimensional feature vector.

To merge features from overlapping windows, we used Gaussian-weighted fusion on the patch-token grid. For a patch location (*i*, *j*) within a window, the fusion weight was defined as

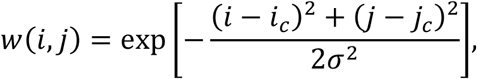

where (*i_c_*, *j_c_*) denotes the center of the 28 × 28 patch grid. We set *σ* = 7, corresponding to one-quarter of the patch-grid side length. For each spatial location in the whole-slide feature map, features contributed by all overlapping windows were accumulated after multiplication by their Gaussian weights and then normalized by the sum of the corresponding weights. This procedure reduces boundary artifacts between adjacent windows and produces a smoothly fused dense feature representation.

The final output for each section *I_i_* is a two-dimensional feature map of size ⌊*H*/16⌋ × ⌊*W*/16⌋ × 768, where *H* and *W* denote the height and width of the input H&E image in pixels. Because all images were resampled to 0.5 μm per pixel, each feature vector corresponds to a 16 × 16-pixel tissue region, equivalent to an approximately 8 × 8 μm² histological unit. To process large serial-section datasets efficiently, feature extraction was parallelized across GPUs, with each GPU worker assigned an independent subset of sections.

#### Morphological feature interpolation

Histological stacks are acquired at finite section intervals and may contain missing or damaged sections, resulting in discrete gaps along the physical *z*-axis. To construct a dense, near-isotropic three-dimensional morphological representation, Spaceland performs optical-flow-based interpolation between adjacent physical sections and synthesizes virtual intermediate histological feature maps at predefined target depths.

For two adjacent physical sections located at depths *z_i_* and *z_i_*_+1_, Spaceland estimates an inter-section displacement field ***u****_i_*_→*i*+1_ using the TV-L1 optical flow algorithm^23,24^ applied to downsampled binary tissue masks. Because the purpose of this step is to guide feature interpolation rather than to perform pixel-level histological registration, the mask-derived flow field was used to capture coarse tissue-shape deformation and boundary displacement between adjacent sections. The estimated displacement field was then upsampled to the feature-map resolution using bilinear interpolation, with displacement vectors scaled according to the downsampling factor. This strategy reduces the computational cost associated with whole-slide images while providing a deformation prior for synthesizing intermediate histological feature maps. TV-L1 optical flow was used to obtain a spatially regularized displacement field from tissue masks while allowing discontinuities near tissue boundaries.

For a virtual section at target depth *z*_target_ between *z_i_* and *z_i_*_+1_, the relative interpolation position was defined as

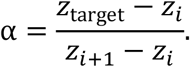

Let *F_i_* and *F_i_*_+1_ denote the histological feature maps extracted from the two adjacent physical sections. A two-sided backward-warping strategy was used to synthesize the virtual feature map *F_v_*(*α*). Specifically, the two adjacent feature maps were warped toward the target depth according to fractional displacements derived from the same inter-section flow field and then combined by distance-weighted linear fusion:

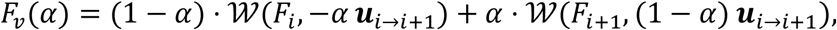

where W(*F*, ***d***) denotes bilinear grid-sampling-based warping of feature map *F* according to displacement field ***d***. Compared with direct linear interpolation between adjacent section features, this deformation-aware interpolation reduces ghosting and structural overlay artifacts caused by residual inter-section misalignment. The interpolated virtual feature maps, together with the physical-section feature maps, form a dense three-dimensional histological feature scaffold for subsequent molecular decoding.

### Sub-spot molecular decoding based on multiple instance learning

To reconstruct gene-expression profiles at sub-spot resolution, Spaceland uses a weakly supervised multiple instance learning (MIL) framework that links high-resolution histological features to low-resolution ST measurements. In this formulation, each ST capture spot provides a pooled molecular label, whereas the histological feature tokens covered by the spot are treated as sub-spot instances. The decoder is trained using spot-level ST supervision and then applied to individual sub-spot instances to infer high-resolution molecular features.

#### Construction of sub-spot instance bags

For each molecularly profiled section, the registered ST spot coordinates were mapped onto the aligned histological feature grid. Based on the physical capture area of the ST platform, such as the 55 μm spot diameter of the Visium platform, all histological feature tokens spatially covered by a capture spot were grouped as a bag of sub-spot instances. For spot *j*, the input bag was represented as

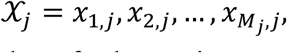

where *M_j_* denotes the number of sub-spot instances within spot *j*, and *x_k_*_,*j*_ ∈ ℝ^d^ is the histological embedding of the*k*-th sub-spot instance. In this study, *d* = 768, corresponding to the feature dimension of the frozen pathology foundation model.

#### Loss function and model optimization

Because ST measurements provide molecular observations at the spot level, model training was performed under weak spot-level supervision. Unlike conventional MIL approaches that use max pooling or attention pooling to select or reweight instances, Spaceland uses summation-based pooling to aggregate sub-spot molecular predictions before comparison with the observed spot-level normalized expression. This design follows the pooled-signal assumption of capture-based ST measurements, in which each spot-level profile represents the combined molecular signal from the tissue area covered by the capture spot.

To make the aggregation consistent with log-normalized expression values, sub-spot predictions were first transformed from log space to linear space, summed within each spot and then transformed back to log space. For gene *g*, the aggregated spot-level prediction was defined as

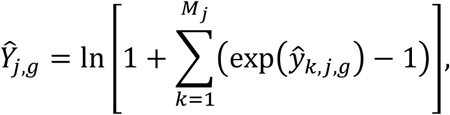

where *ŷ_k,j,g_* is the predicted expression of gene *g* for sub-spot instance *k* in spot *j*, and *Ŷ_j,g_* is the corresponding aggregated spot-level prediction. The training objective was the mean squared error between the aggregated predictions and the observed spot-level expression values:

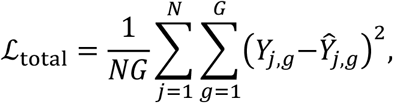

where *N* denotes the number of training spots and *Y_j_*_,*g*_ denotes the observed normalized expression of gene *g* in spot *j*.

The model was implemented in PyTorch Lightning and optimized using the AdamW optimizer with an initial learning rate of 1 × 10^−4^ . Sixteen-bit mixed-precision training was used to reduce memory consumption. Weight decay was applied only to non-bias parameters and parameters outside normalization layers. Model performance was evaluated using five-fold cross-validation, and early stopping based on validation loss was used to reduce overfitting, with a patience of 20 epochs.

### Downstream 3D Analysis

#### 3D expression volume rendering and target gene high-expression domain extraction

To visualize reconstructed three-dimensional gene-expression patterns, consecutive two-dimensional expression maps were converted into physical three-dimensional point clouds. Because high-resolution predicted expression maps contain large numbers of spatial locations, expression matrices were loaded using memory mapping to reduce memory consumption during volume construction. For each section, the two-dimensional in-plane coordinates (*x*, *y*) were combined with the section depth *z*, computed from the section index and physical section thickness, to define a common physical three-dimensional coordinate system.

For each target gene, spatial coordinates and their corresponding expression values were stored as a VTK PolyData object using PyVista and exported as a .vtp file for three-dimensional visualization. Rendering was performed in ParaView by assigning color maps and opacity transfer functions to the gene-expression scalar field.

High-expression domains were extracted using an adaptive gene-specific threshold based on the Kneedle algorithm. For each gene, positive expression values across the reconstructed three-dimensional volume were sorted in descending order. The rank and expression values were normalized to the range [0,1], and the threshold was defined as the expression value corresponding to the point with the maximum perpendicular distance to the diagonal line *x* + *y* − 1 = 0. This geometric knee point identifies the transition between high-expression and low-background regions in the ranked expression distribution. Spatial points with expression values below the gene-specific Kneedle threshold were assigned zero or low opacity, whereas points above the threshold were retained for high-expression domain visualization. This procedure enabled adaptive extraction of target gene high-expression domains in the reconstructed three-dimensional tissue volume.

#### Orthogonal plane virtual slice reconstruction

Using the reconstructed three-dimensional coordinate system and the VTK PolyData representation, Spaceland supports virtual slice reconstruction along orthogonal anatomical planes. For a target axis *a* ∈ *x*, *y*, *z*, a slab region was defined by two parallel clipping planes perpendicular to that axis. The slab was specified by its normal vector, intercept position and physical thickness, thereby selecting spatial points within a defined interval around the target plane.

For each target gene, the selected point-cloud subset and its corresponding expression scalar field were rendered as a two-dimensional virtual slice with finite physical thickness. The slab thickness was chosen according to the spatial resolution of the reconstructed volume and the anatomical scale of the structure being visualized. Color and opacity mappings were applied to the expression values to display molecular patterns within the selected orthogonal plane. This procedure enabled visualization of reconstructed gene-expression distributions along coronal, sagittal and horizontal directions within the same physical coordinate system.

#### Semisupervised 3D spatial anatomical domain identification

Direct unsupervised spatial-domain identification on high-resolution three-dimensional transcriptomic volumes is computationally expensive because of the large number of reconstructed spatial locations. To reduce this cost, Spaceland uses a semisupervised label-propagation strategy. Spatial-domain labels were first obtained by jointly clustering spots from multiple ST-profiled sections using their normalized gene-expression profiles. These cluster assignments were used as spot-level weak labels to train a histology-based domain classifier.

For each profiled section, registered ST spot coordinates were mapped onto the aligned histological feature grid. Each spot was treated as a multiple-instance learning bag, and all 16 × 16-pixel histological feature tiles within its physical capture area were treated as instances. An attention-based multiple instance learning classifier was trained to predict the ST-derived domain label from the corresponding bag of 768-dimensional histological embeddings. The model encoded each tile feature into a 256-dimensional latent representation, computed attention weights across tiles within the same spot, and generated spot-level domain logits through attention-weighted pooling. Training used cross-entropy loss with the ST-derived weak labels, the Adam optimizer and a cosine-annealing learning-rate schedule.

After training, the classifier was applied to the continuous histology-derived morphological scaffold, including both physical-section features and optical-flow-interpolated virtual-section features. During dense inference, each 16 × 16-pixel tile was treated as a singleton instance bag and assigned a domain label. Predictions outside the tissue mask were removed, and the resulting two-dimensional domain maps from physical and virtual sections were assembled in the common three-dimensional coordinate system to obtain high-resolution 3D spatial anatomical domain maps.

### 3D gene co-localization analysis

To quantify spatial co-localization of target genes in the reconstructed three-dimensional expression volume, Spaceland computes a voxel-level gene-set co-localization score from the 3D AnnData expression matrix. For each target gene, expression values across all spatial points were clipped at the 99.5th percentile to reduce the influence of extreme values and low-level background signals, and then rescaled independently to the range [0,1] using min–max normalization. This normalization mitigates the effect of gene-specific differences in expression dynamic range.

For each spatial point, the co-localization score was defined as the product of the normalized expression values of all target genes:

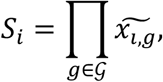

where *G* denotes the target gene set and 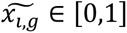 is the normalized expression of gene *g* at spatial point *i*. This product-based score assigns high values only to spatial points where all target genes are jointly elevated, reducing the influence of any single highly expressed gene. The resulting co-localization scores were stored in the observation annotation table of the AnnData object as spatial attributes for downstream three-dimensional rendering, orthogonal virtual sectioning, and axial enrichment analysis.

### Datasets and Biological Setup

#### Mouse Olfactory Bulb (MOB) Dataset

To systematically evaluate Spaceland’s performance under conditions of high axial anisotropy, we constructed a high-density, consecutively sampled mouse olfactory bulb (MOB) dataset. The dataset comprises 360 consecutive coronal sections with 10 μm physical thickness per section. Among these, 180 even-numbered sections were H&E-stained to build a continuous three-dimensional morphological scaffold. Detailed procedures for Decoder-seq spatial transcriptomic profiling, high-resolution ISS validation and ISS image processing are described in **Supplementary Note 2**.

To emulate sparse molecular sampling scenarios typical of large-scale 3D spatial transcriptomics, eight sections (Nos. 40, 80, 120, 160, 202, 240, 280, and 320) were selected for molecular profiling using Decoder-seq spatial transcriptomics. Each ST capture site (spot) covered a square region of approximately 50 μm per side and was used as training supervision for Spaceland.

Additionally, high-resolution in situ sequencing (ISS) data were obtained for three independent sections (Nos. 101, 151, and 201), which were completely held out during training. These sections served as orthogonal benchmarks to rigorously evaluate Spaceland’s ability to predict gene expression at unobserved spatial locations and to assess sub-spot-level reconstruction accuracy.

#### Adult Mouse Hemibrain Dataset

To evaluate the scalability of Spaceland on large tissue volumes with relatively low-resolution spatial transcriptomics, we used the adult mouse hemibrain dataset publicly released by Ortiz et al^32^. This dataset contains sparsely sampled sections along the anterior–posterior axis, each accompanied by paired H&E images, with spatial capture sites approximately 100 μm in diameter. After standard quality control, 62 of the original 75 sections were retained for three-dimensional reconstruction analysis based on anatomical consistency with the mouse hemibrain stereotaxic atlas. Among these sections, 29 were used for model training, while the remaining 33 were reserved as an independent test set to evaluate 3D molecular extrapolation accuracy and spatial reconstruction performance on unseen sections. Detailed anterior–posterior coordinate information for each section is provided in the **Supplementary Table S2**.

#### Planarian Regeneration Spatiotemporal Dataset

To evaluate Spaceland in whole-organism-scale and temporally dynamic tissue reconstruction, we used a multimodal spatiotemporal dataset of *Schmidtea mediterranea* regeneration^34^. The dataset covers six time points spanning intact animals and post-amputation regeneration: intact control, 6 hours post-amputation (6 hpa), 12 hpa, 24 hpa, 3 days post-amputation (3 dpa) and 7 dpa. Each time point contains a series of consecutive H&E-stained tissue sections and paired 10x Genomics Visium spatial transcriptomics data, with Visium capture spots approximately 55 μm in diameter. For each time point, four sections with measured ST profiles were used as molecular supervision for model training. The remaining H&E-only sections were treated as unseen sections for spatial extrapolation and three-dimensional molecular reconstruction. Each time point contained approximately 10–12 consecutive H&E sections, of which 6–8 sections were not used for molecular supervision. Detailed section-level sampling information for each time point is provided in the **Extended Data Fig. 8**.

## Data availability

The in-house mouse olfactory bulb dataset generated in this study will be made publicly available after publication.

## Code availability

The source code for Spaceland, together with documentation and example usage, is being prepared for public release and will be made available in a public repository upon publication.

## Acknowledgements

This work was supported by the National Key Research & Development Program of China (2024YFC3405600 and 2022YFA1104200) to J.S.; and National Natural Science Foundation of China (22474075) to J.S.

## Author contributions

J.S., F.X., Z.Z. and Y.Z. conceived the study. F.X., Z.Z. and Y.Z. performed the major computational analyses, experimental design and data interpretation. F.X. developed and implemented the Spaceland framework. Z.Z. contributed to algorithm development, benchmarking and large-scale data analysis. Y.Z. contributed to experimental data generation, biological interpretation and validation analyses. B.Y., N.H. and W.L. contributed to sample preparation, spatial transcriptomic experiments and data processing. F.X., Z.Z. and Y.Z. generated the figures and visualizations. F.X., Z.Z., Y.Z. and J.S. wrote the manuscript, with input from all authors. J.S., L.W. and C.Y. supervised the project. All authors reviewed and approved the manuscript.

---

**Extended Data Fig. 1.**
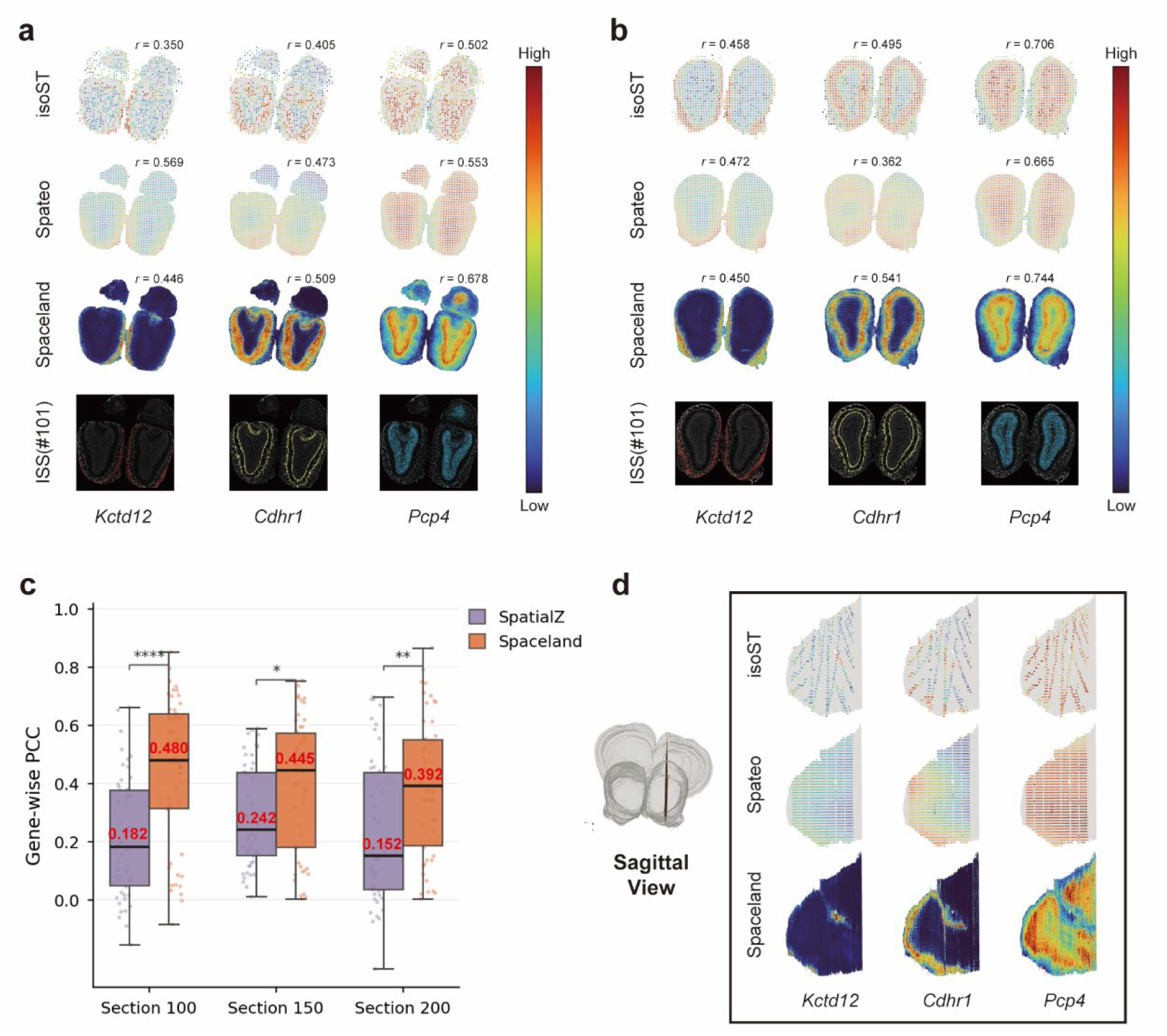
Additional validation of Spaceland reconstruction in unmeasured olfactory bulb sections and 3D molecular fields. a,b, Expression maps of representative laminar markers predicted by isoST, Spateo and Spaceland for molecularly unmeasured sections 150 (**a**) and 200 (**b**), compared with adjacent high-resolution ISS reference sections 151 and 201, respectively. *Kctd12*, *Cdhr1* and *Pcp4* mark the olfactory nerve, mitral cell and granule cell layers, respectively. Pearson correlation coefficients (*r*) between each prediction and the corresponding ISS reference are shown above the maps. **c,** Gene-wise reconstruction accuracy on unmeasured sections 100, 150 and 200. Predictions were evaluated against adjacent ISS references from sections 101, 151 and 201 using the top 50 ISS-derived spatially variable genes ranked by Moran’s I. Each point denotes one gene. Boxes show the median and interquartile range; whiskers extend to 1.5× the interquartile range. Statistical significance was assessed between SpatialZ and Spaceland within each section; *P* values are indicated above the boxes. Red numbers indicate median PCCs. **d,** Sagittal virtual molecular sections queried from reconstructed 3D expression fields. The schematic indicates the queried plane. Expression patterns of *Kctd12*, *Cdhr1* and *Pcp4* are shown for isoST, Spateo and Spaceland.

**Extended Data Fig. 2.**
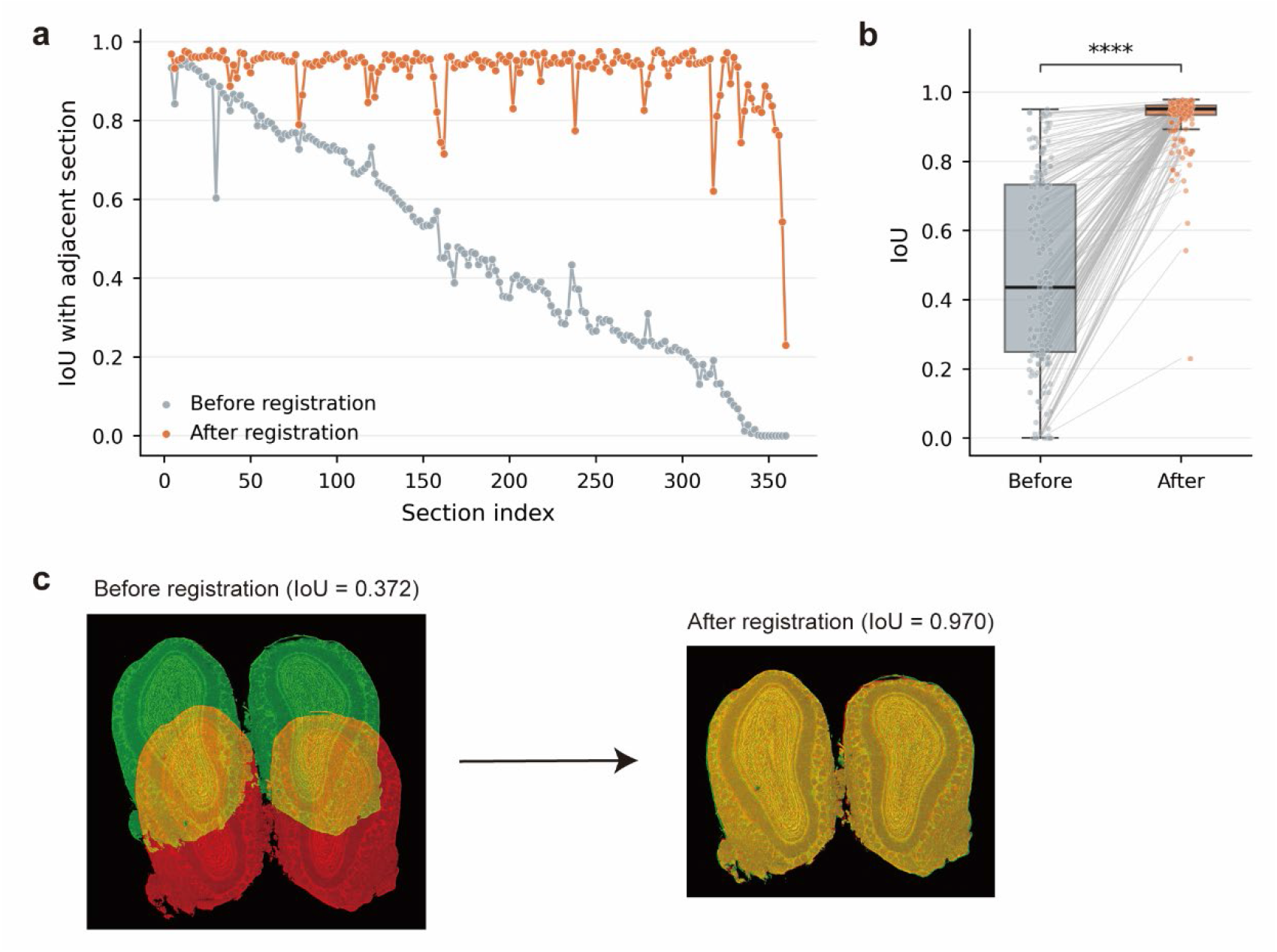
Evaluation of H&E image registration in the mouse olfactory bulb. **a,** Intersection-over-union (IoU) between adjacent H&E sections before and after image registration across the serial mouse olfactory bulb dataset. Registration substantially improved section-to-section anatomical overlap and reduced discontinuities along the section axis. **b,** Paired comparison of adjacent-section IoU values before and after registration. Grey lines connect matched adjacent-section pairs. Box plots show the median, interquartile range and whiskers extending to 1.5× the interquartile range; points denote individual adjacent-section pairs. Statistical significance was assessed using a two-sided paired test; ****P < 0.0001. **c,** Representative overlay of two adjacent H&E tissue masks before and after registration, showing improved anatomical alignment after registration. IoU values for the example pair are indicated above each overlay.

**Extended Data Fig. 3.**
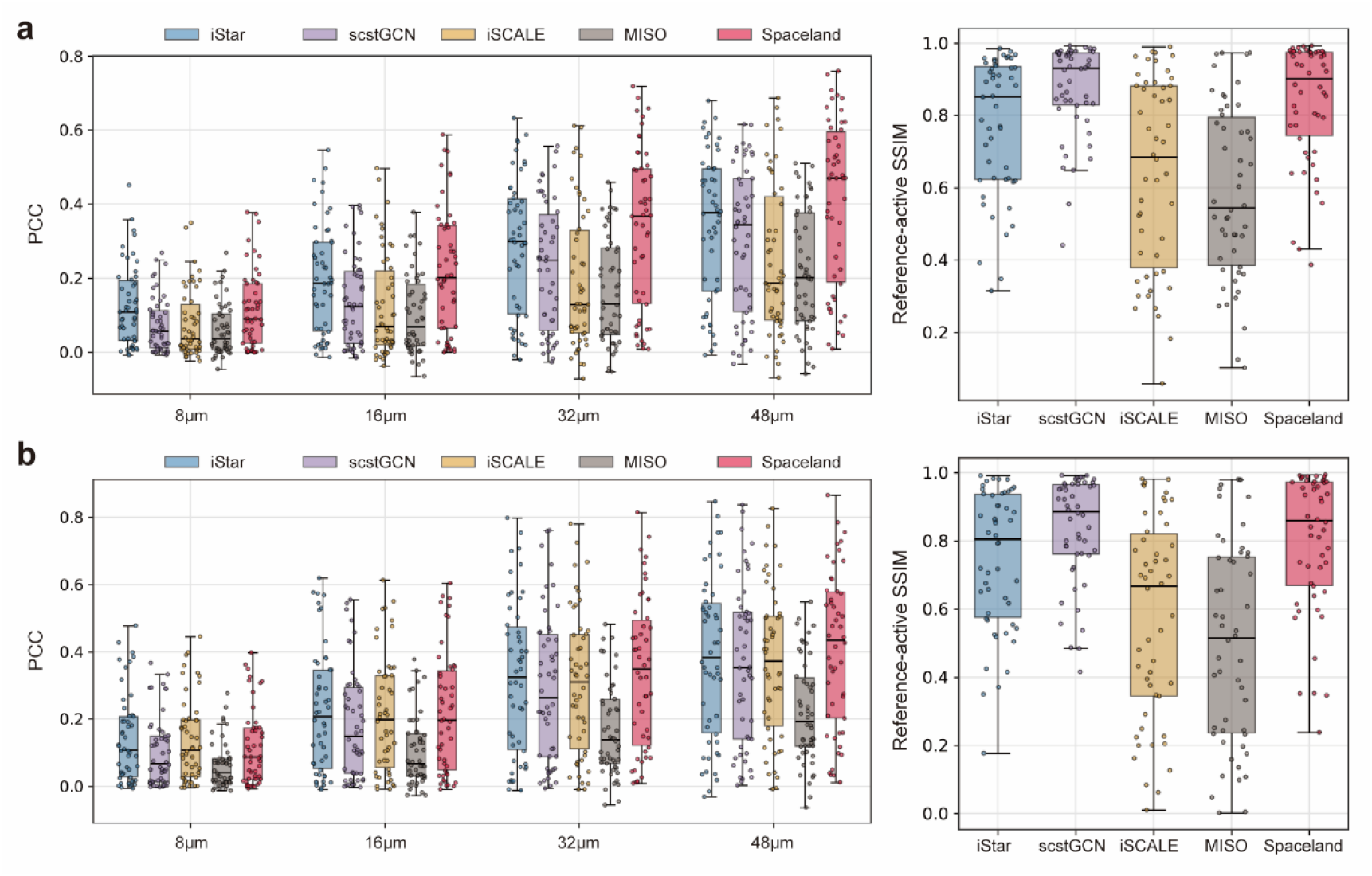
High-resolution orthogonal validation on additional mouse olfactory bulb ISS sections. a,b, Quantitative comparison of Spaceland with representative H&E-based gene-expression prediction baselines on additional mouse olfactory bulb sections. Predictions were generated for molecularly unmeasured H&E sections 150 and 200 and evaluated against adjacent ISS reference sections 151 **(a)** and 201 **(b)**, respectively. For each section, left plots show Pearson correlation coefficients (PCCs) for the top 50 spatially variable genes selected from the ISS reference using Moran’s I, after aggregation to 8, 16, 32 and 48 μm resolutions. Right plots show 8 μm structural similarity index measure (SSIM) computed within GT-active tissue support. For each gene, GT-active pixels were defined from the ISS reference expression map and used to restrict SSIM evaluation to reference-expressing regions, reducing inflation from large shared zero-expression areas. Each point represents one gene at the indicated resolution. Box plots show the median, interquartile range and whiskers extending to 1.5× the interquartile range.

**Extended Data Fig. 4.**
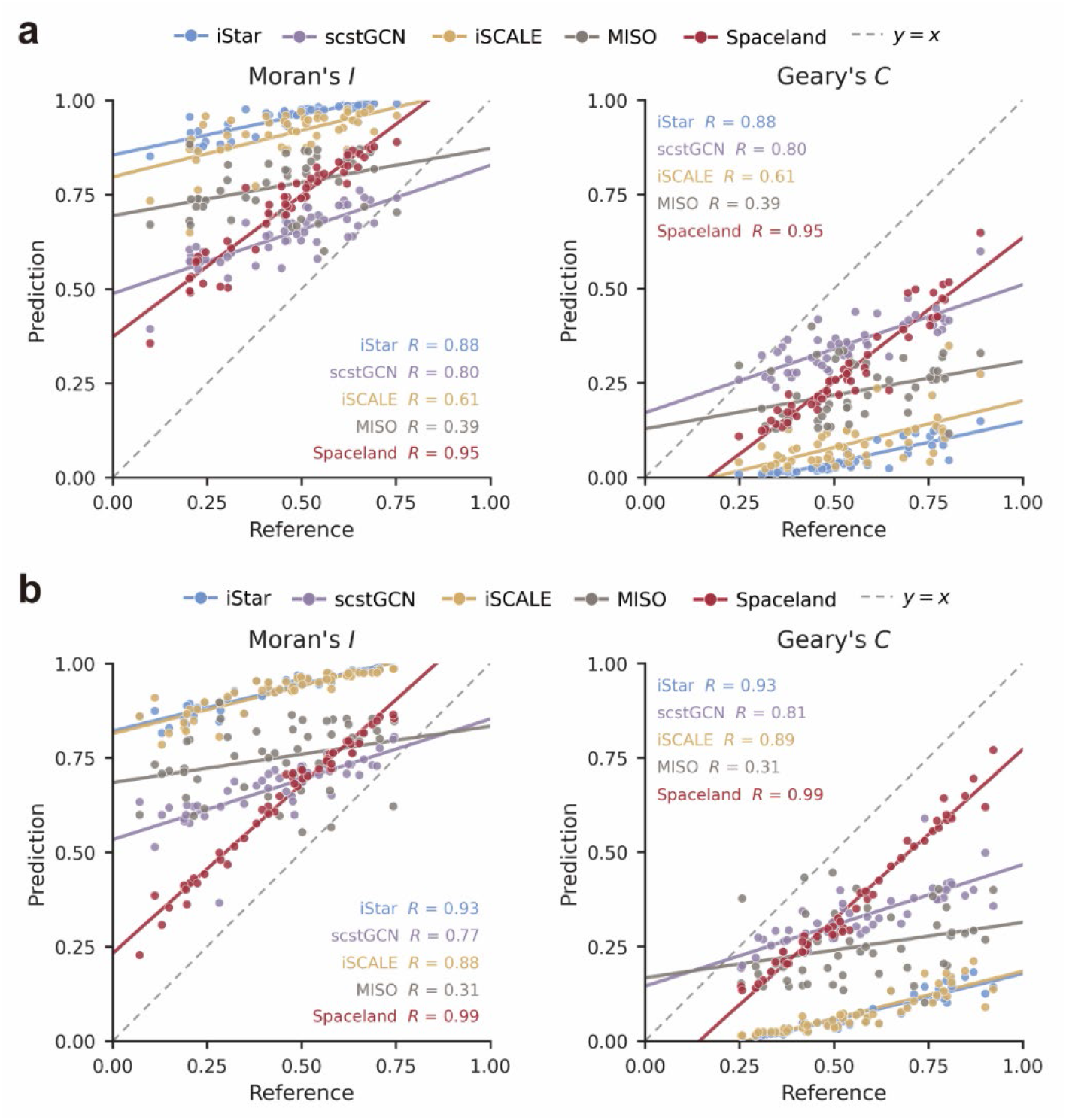
Concordance of spatial autocorrelation metrics between high-resolution predictions and ISS references. a,b, Scatter plots comparing spatial autocorrelation metrics computed from 8 μm high-resolution gene-expression maps predicted by Spaceland and representative H&E-based gene-expression prediction baselines with those computed from adjacent ISS reference sections. Predictions were generated for molecularly unmeasured H&E sections 150 and 200 and compared with ISS reference sections 151 (**a**) and 201 (**b**), respectively. Moran’s I and Geary’s C were calculated for the top 50 spatially variable genes selected from the ISS reference using Moran’s I. Each point represents one gene. Solid lines indicate linear regression fits for each method, and dashed grey lines indicate the identity line, *y* = *x*. Pearson correlation coefficients (*R*) are shown in each panel; all corresponding two-sided Pearson correlation test *P* values were <0.05 and are provided in **Supplementary Data 1**.

**Extended Data Fig. 5.**
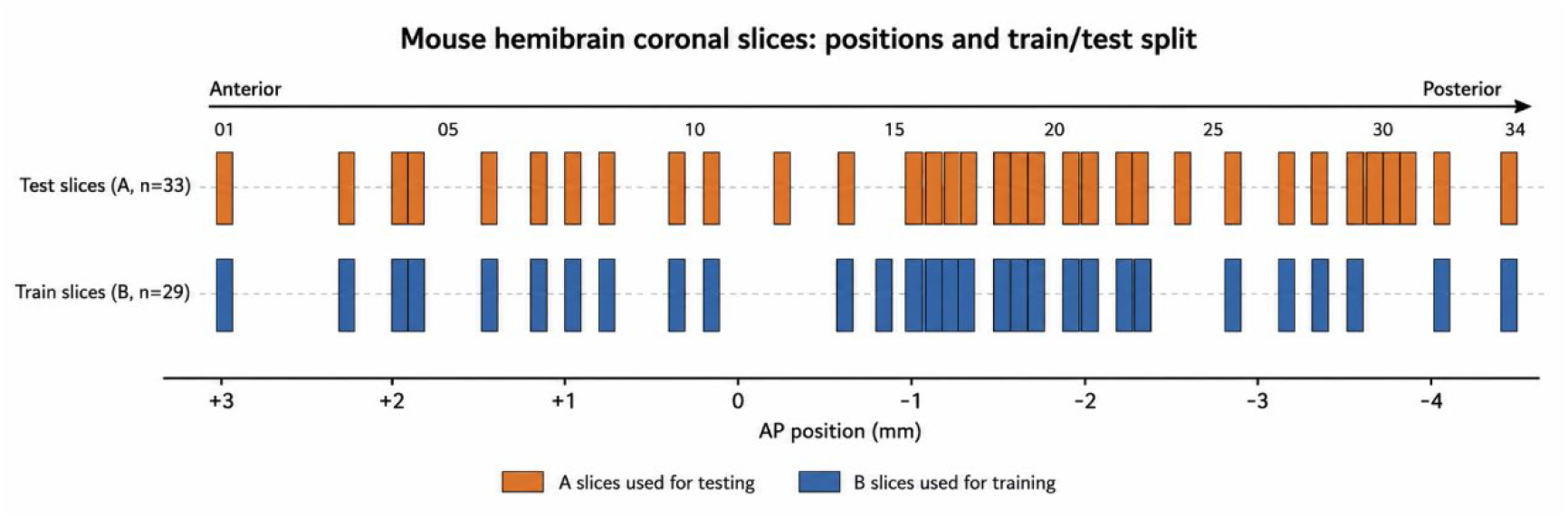
Spatial distribution of training and test sections in the adult mouse hemibrain dataset. Positions of the selected coronal sections along the anterior–posterior (AP) axis used for model training and evaluation. B sections were used for training (n = 29), whereas neighboring A sections were held out for testing (n = 33). Section indices are shown above the axis, and approximate AP positions are indicated below. Detailed AP coordinates and train–test assignments are provided in the **Supplementary Table S2**.

**Extended Data Fig. 6.**
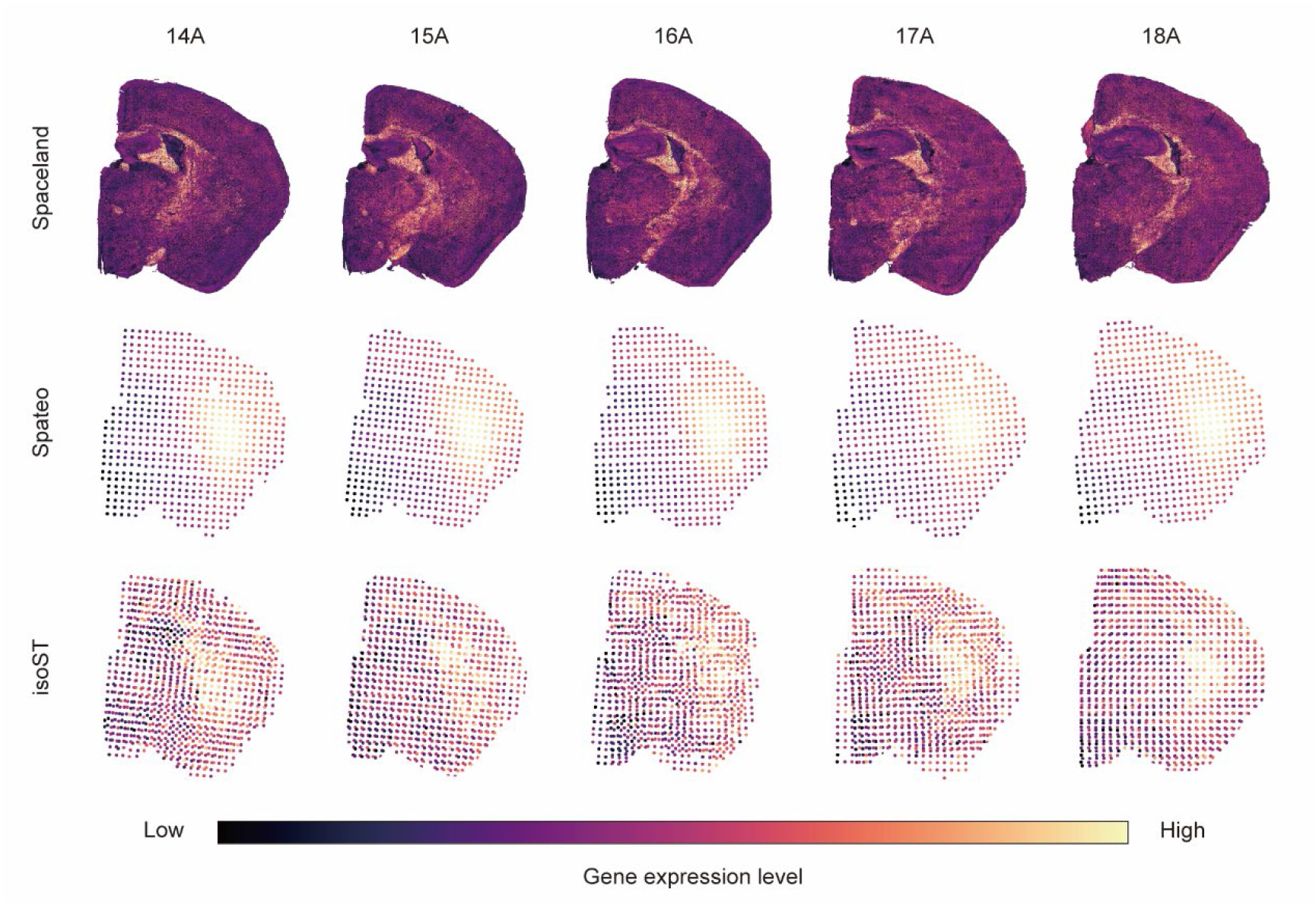
Serial-section visualization of aggregate predicted expression across reconstruction methods. Predicted total expression maps across serial coronal sections 14A–18A are shown for Spaceland, Spateo and isoST. For each section and method, expression values were obtained by summing the predicted expression of 300 genes. Spaceland produces dense tissue-wide expression maps, whereas Spateo and isoST generate spot-level predictions that are shown after rasterization onto the corresponding tissue mask. Color indicates relative aggregate gene-expression level, with darker and brighter colors denoting lower and higher expression, respectively.

**Extended Data Fig. 7.**
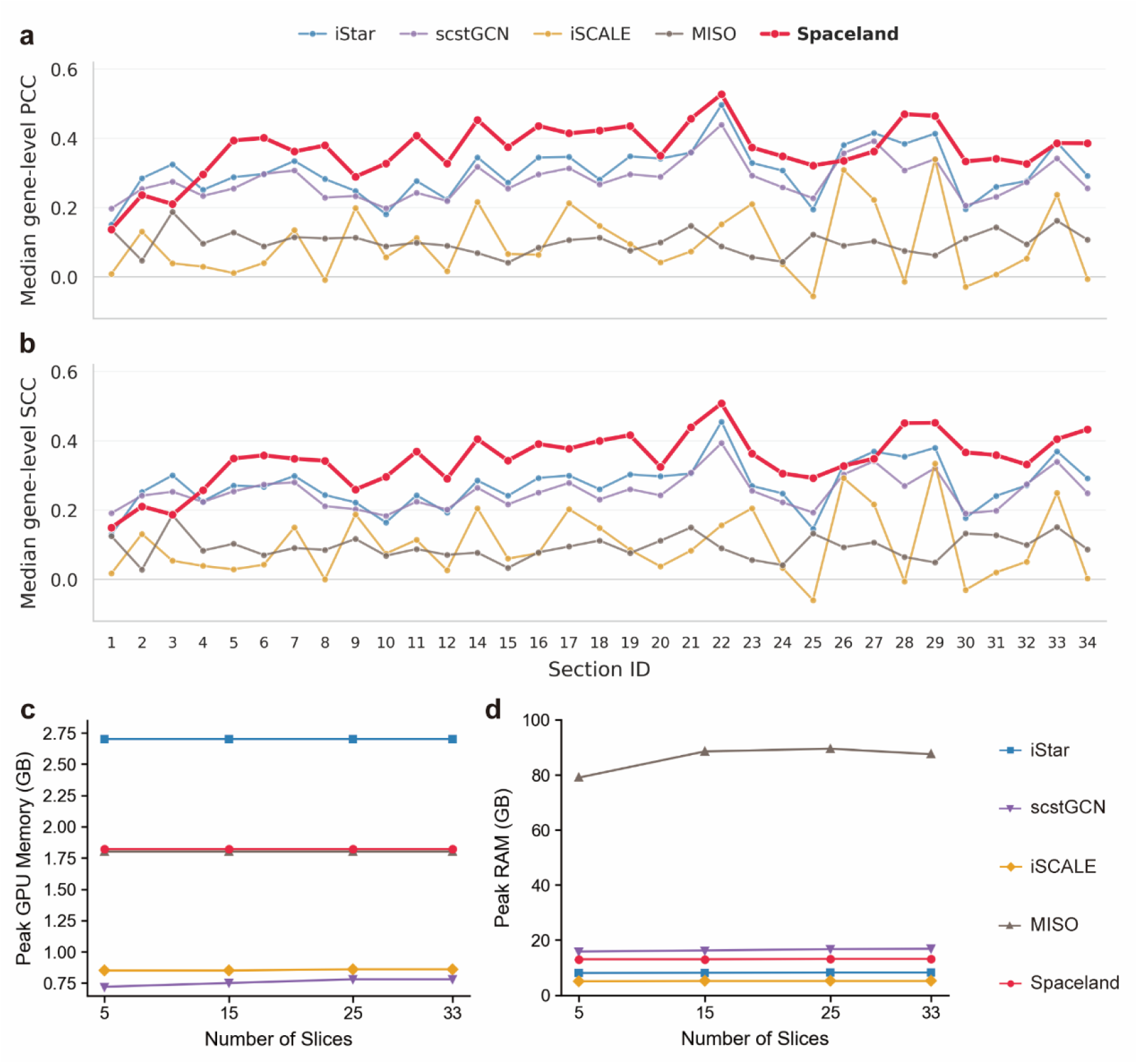
Per-section prediction accuracy and computational profiling in the adult mouse hemibrain benchmark. a,b, Per-section quantitative evaluation of Spaceland and representative H&E-based gene-expression prediction baselines on 33 held-out adult mouse hemibrain sections. For each held-out section, predictions were evaluated across the top 300 spatially variable genes identified using Moran’s I. Lines show the section-wise median gene-level Pearson correlation coefficient (PCC; **a**) and Spearman correlation coefficient (SCC; **b**). **c,d,** Computational resource profiling across reconstruction scales. Peak GPU memory **(c)** and peak system RAM **(d)** were measured while processing increasing numbers of sections. Each method was run independently three times for each setting, and plotted points indicate the mean across runs. All methods were evaluated in the same computational environment.

**Extended Data Fig. 8.**
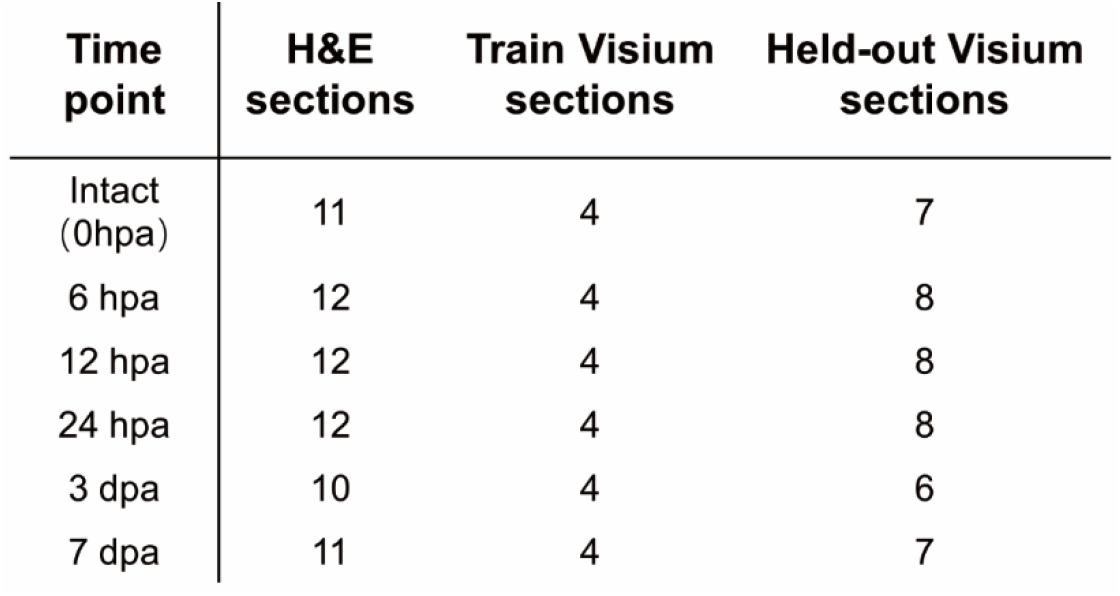
Section composition and train–test splits for the planarian regeneration dataset. Summary of the planarian regeneration dataset used for Spaceland training and evaluation. For each time point, the table lists the number of serial H&E sections, Visium sections used for training and held-out Visium sections used for testing. Four Visium sections were used for training at each time point, with the remaining sections held out for evaluation.

**Extended Data Fig. 9.**
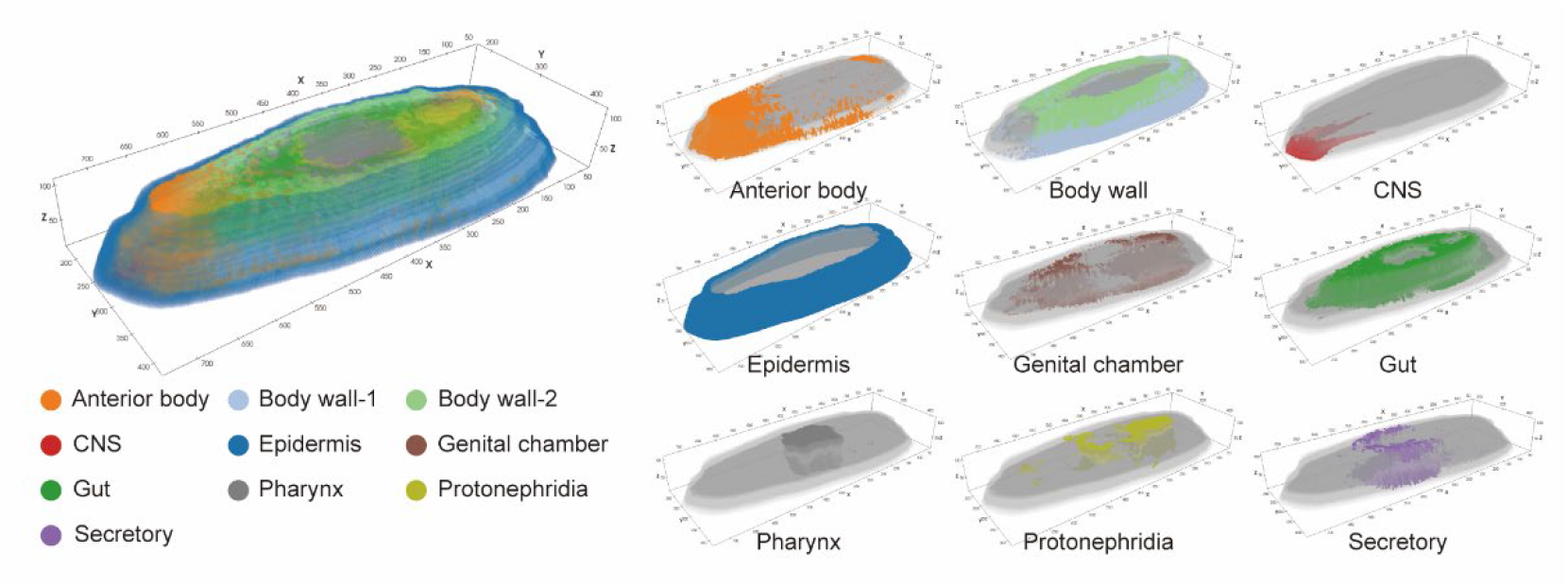
Semi-supervised identification of three-dimensional molecular domains in intact planarians. The left panel shows the integrated domain map, and the right panels show individual domains corresponding to major anatomical structures, including the anterior body, body wall, central nervous system, epidermis, genital chamber, gut, pharynx, protonephridia and secretory tissue.

**Extended Data Fig. 10.**
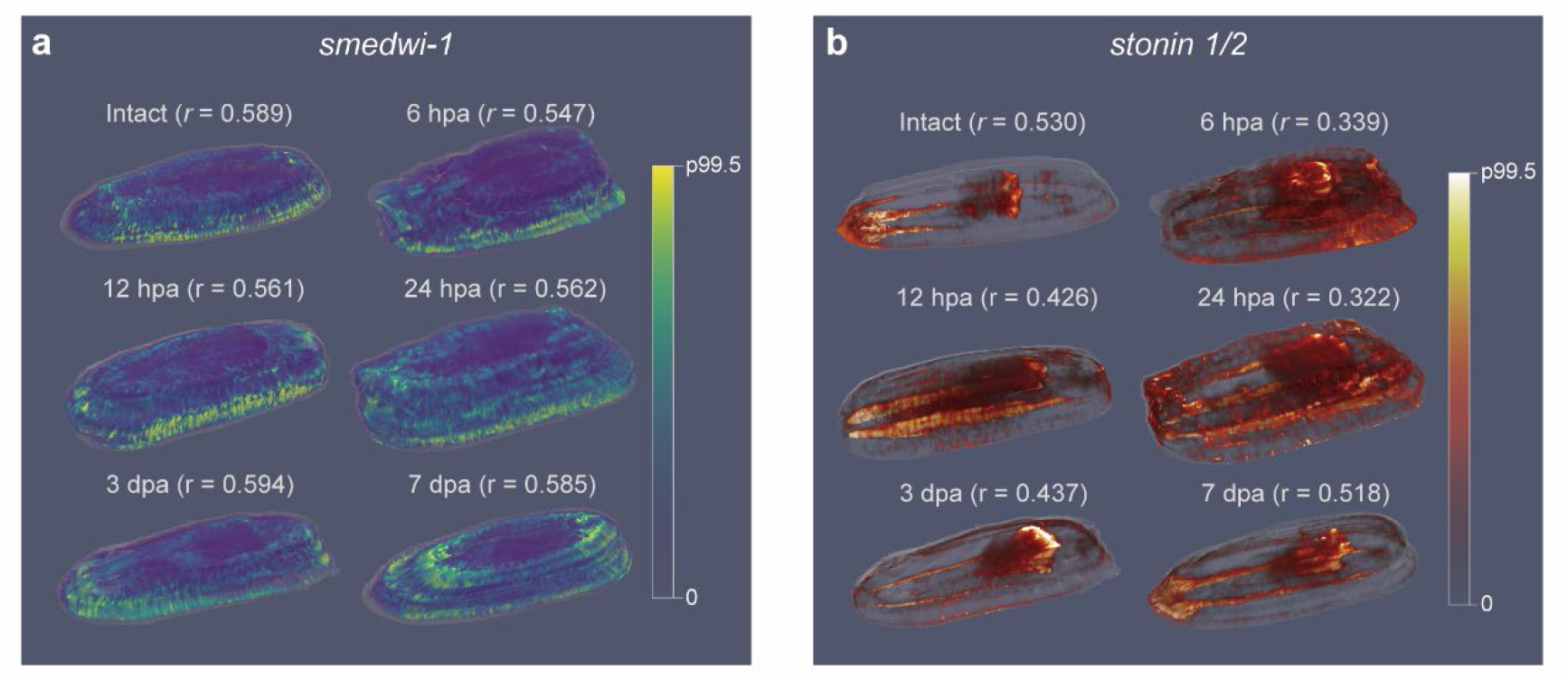
Three-dimensional expression patterns of genes used for planarian co-localization analysis. a,b, Three-dimensional renderings of Spaceland-reconstructed expression patterns for the two genes used in the 3D co-localization analysis during planarian regeneration. Expression patterns are shown for *smedwi-1* (**a**) and *stoning-1/2* (**b**) in intact animals and regenerating animals at 6 hpa, 12 hpa, 24 hpa, 3 dpa and 7 dpa. Pearson correlation coefficients (*r*) between reconstructed and measured expression profiles at each time point are indicated. Expression values are visualized on a 0 to 99.5th percentile scale for each gene.

